# Active DNA demethylation of developmental *cis*-regulatory regions predates vertebrate origins

**DOI:** 10.1101/2021.11.07.467601

**Authors:** Ksenia Skvortsova, Stephanie Bertrand, Danila Voronov, Paul E. Duckett, Samuel E. Ross, Marta Silvia Magri, Ignacio Maeso, Robert J. Weatheritt, Jose Luis Gómez Skarmeta, Maria Ina Arnone, Hector Escriva, Ozren Bogdanovic

## Abstract

DNA methylation (5-methylcytosine; 5mC) is a repressive gene-regulatory mark required for vertebrate embryogenesis. Genomic 5mC is tightly regulated through the coordinated action of DNA methyltransferases, which deposit 5mC, and TET enzymes, which participate in its active removal through the formation of 5-hydroxymethylcytosine (5hmC). TET enzymes are essential for mammalian gastrulation and activation of vertebrate developmental enhancers, however, to date, a clear picture of 5hmC function, abundance, and genomic distribution in non-vertebrate lineages is lacking. By employing base-resolution 5mC and 5hmC quantification during sea urchin and lancelet embryogenesis, we shed light on the roles of non-vertebrate 5hmC and TET enzymes. We find that these invertebrate deuterostomes employ TET enzymes for targeted demethylation of regulatory regions associated with developmental genes and show that the complement of identified 5hmC-regulated genes is conserved to vertebrates. This work thus demonstrates that active 5mC removal from regulatory regions is a common feature of deuterostome embryogenesis suggestive of unexpected deep conservation of a major gene-regulatory module.

## Introduction

Metazoan development is governed by the activity of *cis*-regulatory DNA elements that serve as binding platforms for transcription factors controlling spatial and temporal gene expression^1-4^. In vertebrates, one of the major gene-regulatory mechanisms entails methylation of the 5^th^ carbon of the cytosine pyrimidine ring (5mC) within the DNA^5^. 5mC is a predominantly repressive gene-regulatory mark, which in vertebrates displays unparalleled developmental dynamics. Mammalian embryogenesis is characterised by two rounds of genome-wide 5mC reprogramming that take place in the preimplantation embryo and in the developing germline^6,7^. Global 5mC erasure occurs through a combination of active and passive 5mC removal mechanisms, such as the exclusion of maintenance DNA methyltransferase (DNMT1) from the nucleus^8-12^, and targeted oxidation of 5mC by Ten Eleven Translocation (TET) enzymes, respectively^13-15^. TET enzymes oxidise 5mC through a series of reactions involving 5-hydroxymethylcytosine (5hmC), 5-formylcytosine (5fC), and 5-carboxylcytosine (5caC) intermediates, the latter two of which can be excised by base excision repair (BER) pathways, thus leading to the restoration of unmethylated cytosine at affected sites^16,17^. Apart from participating in mammalian global 5mC erasure, TET enzymes drive targeted 5mC removal from promoters and enhancers required for vertebrate gastrulation^18^ and organ formation^19^. TET activity is thus crucial for orchestrating gene regulatory programs associated with cell fate determination and tissue differentiation. TET depletion accompanied by 5hmC reduction results in enhancer hypermethylation and reduced chromatin accessibility in vertebrate embryos^19,20^, delayed gene induction during cellular differentiation^21^, and skewed lineage differentiation^22-24^. Loss of 5mC in DNMT mutant cells, on the other hand, was shown to result in the activation of intragenic enhancers^25^, altogether highlighting the significance of timely DNA demethylation in the regulation of *cis*-regulatory elements and lineage specification.

In stark contrast to vertebrates, which are characterized by high genomic 5mC levels (∼80%), most nonvertebrate organisms sampled to date possess mosaic DNA methylation patterns that vary widely in their content, genomic distribution, and function^26-28^. Notable exceptions to that rule include *Drosophila melanogaster*^29^ and *Caenorhabditis elegans*^30^ whose genomes are devoid of 5mC, and the demosponge *Amphimedon queenslandica*, the genome of which displays vertebrate-like genomic hypermethylation^31^. Those exceptions notwithstanding, a common feature of invertebrate 5mC patterning is the association of 5mC with gene bodies of actively transcribed genes^26^. 5mC has been shown to prevent precocious transcription from cryptic transcription start sites located within gene bodies^32^. However, it is currently underexplored whether any other aspects of 5mC-mediated genome regulation are evolutionarily maintained across the invertebrate-vertebrate boundary. For example, TET proteins are present in most non-vertebrate genomes^33^ where their function has been linked to RNA hydroxymethylation^34^, production of 5hmC in the germline^35^, or regulation of alternative splicing^36^. Except for a single study that employed ELISA-based approaches to demonstrate global 5hmC developmental loss in the Pacific sea gooseberry (*Pleurobrachia bachei*)^37^, it is not yet clear to what extent non-vertebrate organisms utilise 5mC and consequently TET proteins and 5hmC for developmental gene regulation. Moreover, base-resolution hydroxymethylomes have thus far only been generated for sponges and sea anemones (discussed below)^31^, leaving numerous open questions as to the quantity, genomic localisation, and function of non-vertebrate 5hmC.

While it is well-established that non-vertebrate genomes exhibit low 5mC, it is worth noting that *cis*-regulatory elements frequently reside within highly methylated gene bodies. It is thus possible that the 5mC state of such elements is regulated independenttly of bulk gene body 5mC. For example, we have recently shown that the European lancelet (*Branchiostoma lanceolatum*), an invertebrate chordate, exhibits developmental DNA demethylation of *cis*-regulatory elements located within methylated gene bodies^38^. Notably, this demethylation event coincided with the expression of the lancelet TET orthologue, thereby suggesting that animals with mosaic 5mC might employ TET-mediated demethylation of enhancers for their spatio-temporal regulation during development. 5hmC profiling of genomic DNA extracted from the demosponge *A. queenslandica* demonstrated enrichment of this mark at potential transcription factor binding sites^31^. Nevertheless, definitive proof that TET proteins and active DNA demethylation participate in developmental enhancer or promoter usage in nonvertebrate lineages, has not been provided to date. To better understand the evolution of TET-mediated regulation of distal regulatory elements, we conducted detailed analyses of genome-wide 5mC and 5hmC dynamics during development of the purple sea urchin (*Strongylocentrotus purpuratus*), which forms part of the sister lineage to chordates, the ambulacrarians^39^, and the invertebrate chordate European lancelet^38^. Specifically, we employed whole genome bisulfite sequencing (WGBS)^40^ and APOBEC-coupled epigenetic sequencing (ACE-seq)^41^ to generate base-resolution maps of 5mC and 5hmC, respectively, at four developmental time points. Integration of these datasets with open chromatin profiles demonstrated prominent active DNA demethylation of regulatory regions in both examined organisms, thus strongly supporting the notion that invertebrate deuterostomes employ TET proteins and 5hmC for developmental gene regulation. Moreover, we found that vertebrate orthologues of genes marked by 5hmC in these invertebrate deuterostomes are associated with 5hmC enrichment and defined spatio-temporal regulatory logic in zebrafish. Altogether, our work demonstrates that at least in some non-vertebrate lineages, 5mC is not merely a by-product of geneactivity required for the prevention of spurious transcriptional initiation, but a *bona-fide* regulatory mark that is being actively remodelled at regulatory regions associated with key developmental processes.

## Results

### Structural and functional conservation of TET enzymes

To better understand the evolutionary conservation of 5hmC and TET expression dynamics during embryogenesis, we first wanted to determine whether echinoderm and cephalochordate TET enzymes are characterised by the same protein domain composition as vertebrate TETs. The sea urchin genome^39^ as well as the genomes of the European^38^ and Florida lancelet^42^ encode a single copy of TET proteins. The genomes of most vertebrates, however, exhibit two or three TET copies^43,44^ (**Fig. S1A**), which arose during the process of whole genome duplication^45^. Comparison of TET protein structure revealed that both the sea urchin TET (sTET) as well as the lancelet TET (bTET) contain an N-terminal CxxC domain, which is also present in vertebrate TET1 and TET3 enzymes (**Fig. 1A**)^46^. CxxC domains can interact directly with the DNA thereby facilitating TET interaction with chromatin^47,48^. The C-terminus of sTET and bTET is characterised by a doublestranded β-helix (DSBH) domain that harbours binding sites for the Fe(II) and 2-oxoglutarate (2-OG) cofactors required for TET enzymatic activity^46^. Notably, the DSBH domain in sTET, bTET and vertebrate TET proteins is interrupted by a sizable low complexity region (**Fig. 1A**). These results are in line with the previously described structural conservation of metazoan TET proteins^33^. We next conducted multiple sequence alignment analyses and found strong sequence conservation in the core catalytic domain (**Fig. 1B, Fig. S1B**), as well as striking structural conservation between sTET, bTET and their vertebrate counterparts (**Fig. 1C, Fig. S1C-E**). Having determined the suitability of sea urchin and lancelet for TET evolutionary comparisons and 5hmC profiling, we next quantified TET expression during sea urchin, lancelet, and zebrafish development^38,49^. In all examined species, the developmental peak of TET expression could be observed during mid-development (**Fig. 1D**), coinciding with gastrulation (sea urchin) and segmentation (lancelet, zebrafish) (**Fig. S1F**)^50,51^. These observations are in accord with reduced developmental requirements for TET activity during vertebrate pluripotency^18-20,22^. Overall, our results demonstrate highly conserved TET protein structure and developmental TET expression profiles across diverse deuterostomes, suggestive of the important roles these proteins might play during animal development.

**Figure 1.**
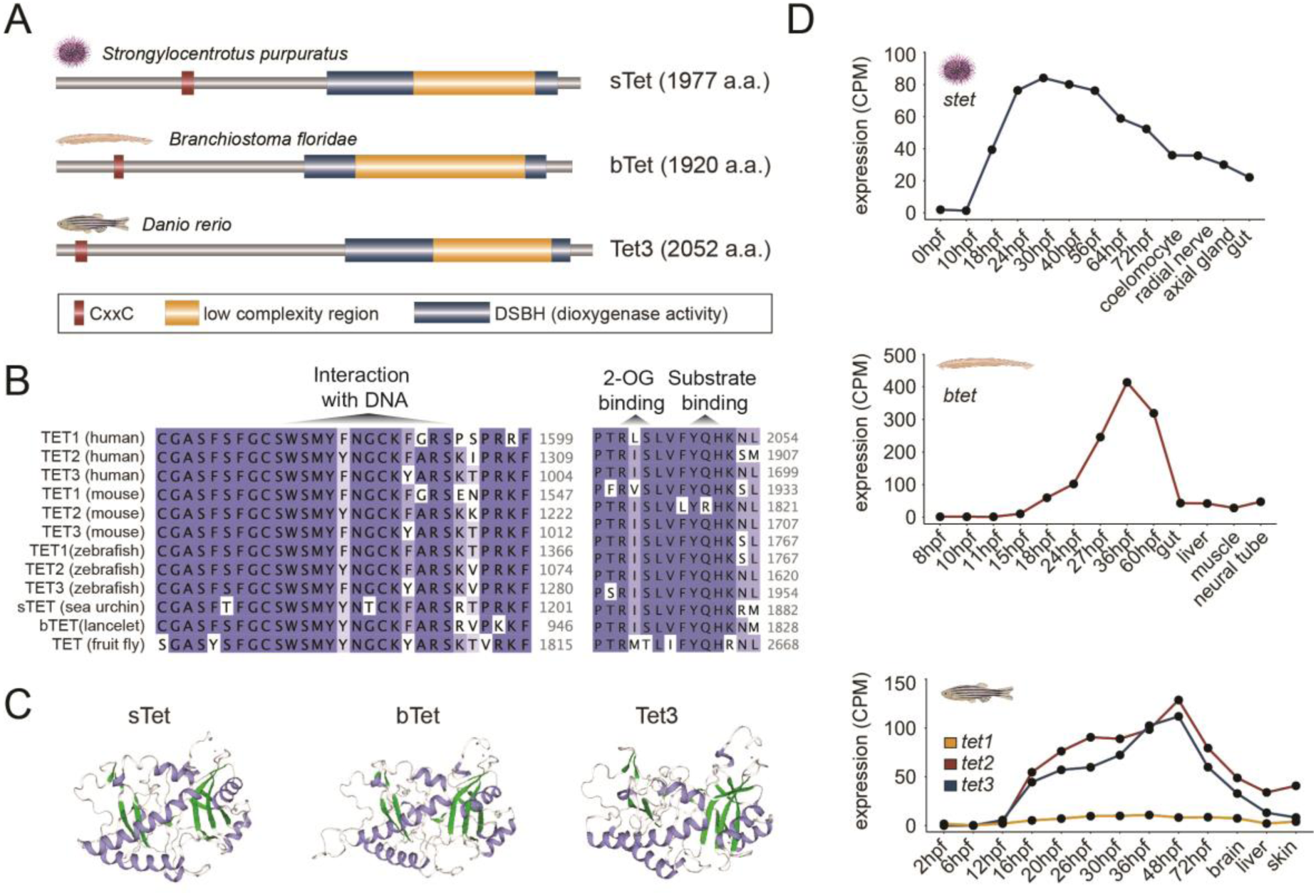
Evolutionary conservation of TET protein sequence, catalytic domain structure, and developmental expression. **(A)** Sea urchin, lancelet and zebrafish TET protein domain structure predicted using SWISS-MODEL and HMMER. Sea urchin sTet, Florida lancelet bTet, and zebrafish Tet3 harbour an N-terminal DNA-binding CxxC domain and a C-terminal catalytic double stranded β-helix domain (DSBH) containing a large low-complexity insert. **(B)** Multiple sequence alignment of human, mouse, and zebrafish TET1, TET2, TET3 as well as sea urchin, lancelet, and fruit fly TET core catalytic domain. A part of DSBH is shown. The colour of each amino acid indicates percentage identity (PID) with darker blue depicting higher PID and lighter blue - lower PID. **(C)** 3D models of methylcytosine dioxygenase domains of sea urchin sTet, lancelet bTet, and zebrafish Tet3 performed using SWISS-MODEL. Crystal structure of the human TET2-5fC complex (5d9y.1.A) was ranked as a top template in the template search and was used for model building. α-helix is depicted in blue and β-sheet - in green. **(D)** TET gene expression dynamics during sea urchin, lancelet, and zebrafish development.

### DNA methylation content and dynamics of sea urchin and lancelet genomes

To quantify the amount of genomic 5mC, which can act as substrate for TET oxidation and 5hmC formation, we generated base-resolution 5mC profiles (WGBS)^40^ corresponding to the following stages of sea urchin development: 24hpf (blastula), 48hpf (gastrula), 72hpf (pluteus), adult (tube feet). Additionally, we analysed WGBS datasets of lancelet embryogenesis from 8hpf (blastula - G3), 15hpf (gastrula - G6), 36hpf (neurula - T0), and adult (liver - A) time points (**Fig. 2A, Table S1**)^38,50,51^. In agreement with previous studies^38,52^, average methylation values (sea urchin = 23-28%, lancelet = 21-25%) (**Fig. 2A)** were consistent with canonical invertebrate methylome patterns, where the majority of genomic 5mC is located within gene bodies of expressed genes (**Fig. S2A-C**)^28^. Notably, the adult samples of both the sea urchin and the lancelet displayed considerably lower 5mC levels reminiscent of developmental 5mC dynamics in zebrafish and *Xenopus tropicalis* (**Fig. 2A)**^19^. Given the positive correlation between gene expression and gene body 5mC in non-vertebrate organisms, lower 5mC levels in adult tissues could potentially be explained by specialised transcriptional profiles of those tissues characterised by a smaller number of expressed genes or a lower transcriptional output overall. To that end, we analysed the transcripttomes^38,49^ of multiple developmental stages and adult tissues and revealed comparable mean expression profiles across all examined samples in both organisms (**Fig. 2B)**. These results suggest that the observed loss of 5mC in adults is likely not caused by major changes in transcriptional complexity but is rather linked to a gradual increase in chromatin accessibility associated with more frequent usage of *cis*-regulatory elements^53^. To obtain further insight into developmental dynamics of 5mC, we identified genomic regions, which display significant (*FDR* < 0.05, ΔmCG/CG ≥ 0.2) developmental changes in 5mC levels (**Fig. 2C, Table S2-S7**).

**Figure 2.**
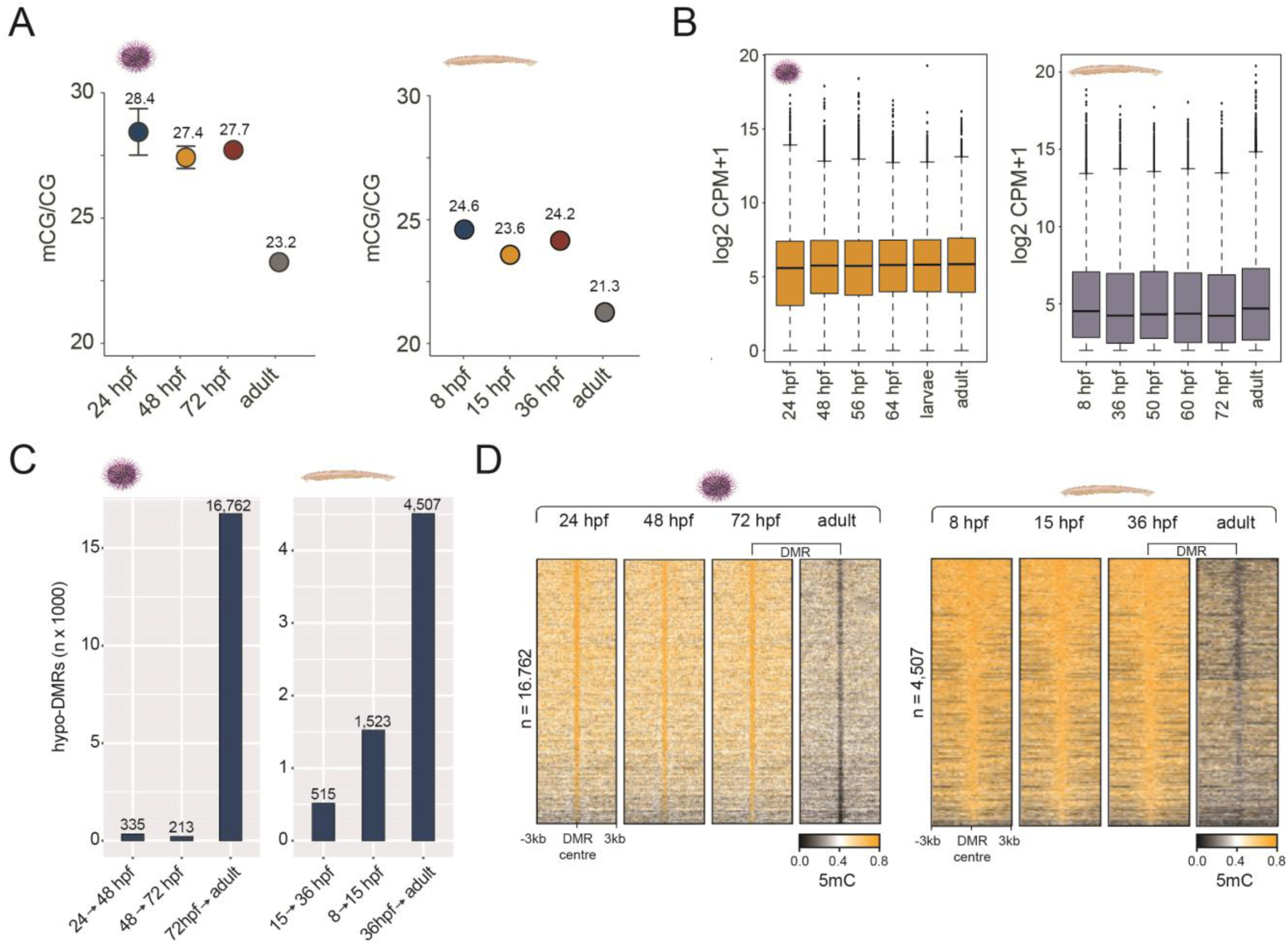
DNA methylation dynamics of sea urchin and lancelet development. **(A)** Average DNA methylation (mCG/CG) levels during sea urchin and lancelet development. **(B)** Average gene expression (CPM+1) levels during sea urchin and lancelet development. **(C)** Number of hypomethylated DMRs (hypoDMRs) identified between embryonic and adult stages in sea urchin and lancelet genomes. **(D)** Heatmaps depicting DNA methylation (mCG/CG) at adult hypoDMRs in sea urchin and lancelet genomes.

We identified 20,072 and 8,136 of such differentially methylated regions (DMRs) in sea urchin and lancelet genomes, respectively, with the majority corresponding to the larva/pluteus-to-adult transitions (**Fig. 2D, Fig. S2D**). The majority (86% sea urchin, 80% lancelet) of the identified DMRs were associated with developmental hypomethylation (**Fig. 2C-D, Fig. S2D**). These results are in agreement with the overall lower levels of 5mC in adult tissues in both organisms (**Fig. 2A)**. Altogether, we find that the embryogenesis of sampled invertebrate deuterostomes is characterised by significant developmental loss of 5mC, which cannot be explained by developmental changes in transcriptional activity.

### Developmental dynamics of the 5-hydroxymethylcytosine landscape

To explore the developmental dynamics, genomic content, and distribution of 5hmC in sea urchin and lancelet genomes and to compare those against vertebrate 5hmC patterns, we employed nondestructive, base-resolution sequencing of 5hmC using a DNA deaminase (ACE-seq)^41^. We first wanted to test how ACE-seq 5hmC quantification compares to TET-assisted bisulfite sequencing (TAB-seq)^54^ and immunoprecipitation (hMeDIP-seq) ^55^ 5hmC profiles in zebrafish, an organism with a well-described DNA hydroxymethylome^19,22,56,57^. Firstly, we found that the same genomic regions displayed 5hmC enrichment identified by all the three techniques (**Fig. S3A, B**). Furthermore, we observed a significant overlap between data originating from the two base-resolution approaches (**Fig. S3C**) and hence proceeded to generate sea urchin and lancelet 5hmC maps using ACE-seq. We obtained DNA hydroxymethylome profiles for the following stages of sea urchin and lancelet embryogenesis: 24hpf (blastula), 48hpf (gastrula), 72hpf (pluteus), adult (tube feet), and 36hpf (late neurula - T0), 60-72hpf (larvae - L0), 3-4mo (juvenile - J), adult (liver - A)^50,51^ **(Table S1**). Identification of hydroxymethylated sites (proportion test, *P* adj. < 0.05, ΔhmCG/CG ≥ 0.1)^58^ in sea urchin and lancelet genomes, revealed 1-2% of CpGs with significant 5hmC content (**Fig. 3A)**. In zebrafish 24hpf embryos, which are characterised by strong TET activity (**Fig. 1D**), ∼5% of overall CpGs were identified as significantly hydroxymethylated (**Fig. 3A)**. These results are in line with the overall higher global 5mC content of vertebrate genomes (**Fig. 3A)**. To assess the reproducibility of ACE-seq 5hmC data between replicates, we searched the genome for sites of significant 5hmC enrichment and plotted mean 5hmC values between replicates (**Fig. 3B**). We observed an overall positive correlation of 5hmC levels in those regions with correlation values of *R*=0.58 and *R*=0.68 for sea urchin and lancelet, respectively. Given the transient nature of 5hmC and its role in 5mC to C conversion, the expectation would be for the majority of 5hmC sites to exist in a fully methylated (5mC) state before the onset of TET activity and in a fully demethylated (C) state once the period of TET-mediated demethylation has ended. To test whether this holds true in amphioxus and sea urchin genomes, we identified significant 5hmC sites at the stage coinciding with high TET expression (48hpf – sea urchin; 60hpf lancelet) and interrogated their fate before and after this period (**Fig. 3C, D**). Expectedly, we found that the majority of hydroxy-methylated CpGs were in the 5mC state before the mid-developmental period. The same fraction also displayed a notable enrichment in unmethylated cytosines in adult tissues, in agreement with developmental 5mC removal. These data thus demonstrate that ACE-seq can accurately detect 5hmC at base resolution in sea urchin and lancelet genomes and that 5hmC displays comparable developmental dynamics across deuterostomes.

**Figure 3.**
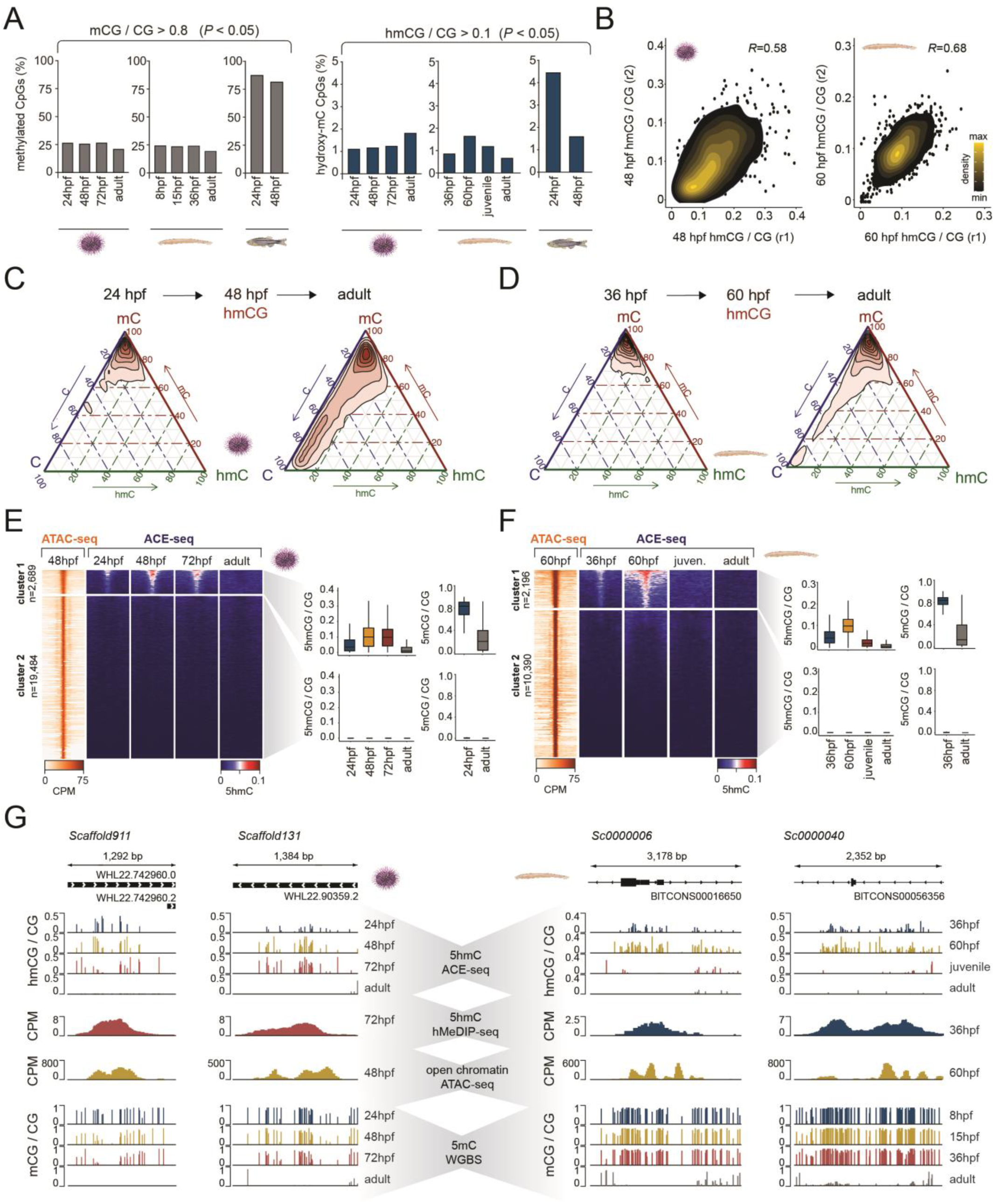
DNA hydroxymethylation landscape of sea urchin and lancelet development. **(A)** Percentage of methylated and hydroxymethylated CpGs in embryonic and adult tissues of sea urchin, lancelet, and zebrafish. CpG sites with at least 80% methylation and CpG sites with at least 10% hydroxymethylation (*P* adj. < 0.05) were included in the analysis. Only CpG sites with a minimum of 10x coverage were considered. **(B)** Concordance of hydroxymethylation levels between biological replicates. Sea urchin 48hpf and lancelet 60hpf samples are shown. **(C, D)** Ternary plots depicting the relationship between methylated (mC), hydroxymethylated (hmC) and unmethylated (C) CpG states in sea urchin **(C)** and lancelet **(D)** embryos and adult tissues. CpG sites hydroxymethylated at 48hpf in sea urchin **(C)** and 60 hpf in lancelet **(D)** (*P* adj. < 0.05, proportion test), are shown. **(E, F)** Heatmaps depicting DNA hydroxymethylation dynamics (ACE-seq) of open chromatin regions (ATAC-seq) in sea urchin **(E)** and lancelet **(F)**. K-means clustering (k=2) of ACE-seq and ATAC-seq signal over ATAC-seq peaks. Boxplots of DNA hydroxymethylation and DNA methylation dynamics during sea urchin **(E)** and lancelet **(F)** development. **(G)** IGV browser tracks depicting DNA hydroxymethylation (ACE-seq and hMeDIP-seq) enrichment at open chromatin regions (ATAC-seq) coinciding with DNA demethylation (MethylC-seq) in sea urchin and lancelet genomes.

### 5-hydroxymethylcytosine marks regulatory regions in deuterostome genomes

Next, we wanted to assess to what extent does 5hmC correlate with sites of developmental hypomethylation and regulatory regions in general, identified through open chromatin profiling. We first plotted the 5hmC signal over previously identified adult hypomethylated DMRs (**Fig. 2**) and found a notable overlap between the two datasets (**Fig. S3D**). Out of 16,753 DMRs, 5,585 (33%) formed an 5hmC-enriched cluster in the sea urchin, whereas in the lancelet that overlap was 46% (2,072 out of 4,507 DMRs). To obtain more comprehensive insight into the relationships between 5hmC and gene regulation, we generated open chromatin profiles (ATAC-seq), corresponding to 48hpf (sea urchin) and 60hpf (lancelet)^38^ stages (**Fig. 3E, F**), which are associated with peaks of TET activity (**Fig. 1D**). K-means clustering (k=2) of 5hmC signal over those regions revealed a subset (cluster 1), which was enriched for 5hmC (**Fig. 3E, F**). This cluster was characterised by high 5mC levels in embryonic and low 5mC levels in adult tissues, in agreement with active demethylation occurring at these regions. 5hmC was highly enriched in 48hpf and 60hpf embryos in sea urchin and lancelet, respectively, and was almost undetectable in adult samples, as expected from the low 5mC signal in those tissues (**Fig. 3E-G, Fig. S3E, F**). Altogether, these results demonstrate strong 5hmC presence at invertebrate regulatory regions, which coincides with the active removal of 5mC.

### 5-hydroxymethylcytosine and developmental gene activation

Having characterised developmental dynamics of 5hmC (**Fig. 3**), we next sought to understand which gene groups are potentially regulated by 5hmC and what impact might 5hmC have on developmental gene regulation. We first generated gene ontology (GO) profiles^59^ of genes associated with 5hmC-marked ATAC-seq peaks^60^ and found significant enrichment for developmental processes (**Fig. 4A**). Many of these processes such as “system development”, “cell differentiation”, “tissue development”, “developmental process”, and others, were identical to those associated with 5hmC-regulated genes previously described in zebrafish, *Xenopus*, and mouse embryos^19^. We then assessed the genomic distribution of 5hmC enrichment and found that the majority of 5hmC peaks reside in gene bodies, even though a significant fraction of 5hmC was observed in promoters and intergenic regions (**Fig. 4B**). In line with their proposed developmental function, genes with both promoter and gene body 5hmC peaks were on average longer, suggestive of their increased regulatory potential **(Fig. 4C, Fig. S4A, B)**^61^. Transcription factor binding motif analyses revealed significant enrichment for Sox and Fox binding sites, both of which have been previously shown to associate with TET proteins and DNA demethylation in vertebrates^62,63^ (**Fig. 4D, Fig. S4C, D, Tables S8-13**). In vertebrates, promoter 5hmC is linked to transcriptional activation^64,65^. To test whether a similar association could be reproduced in sea urchin and lancelet genomes, we analysed the transcriptional profiles of genes with a promoter 5hmC peak and found that promoter 5hmC correlates with activation of developmental genes in both examined organisms (**Fig. 4E**). A similar positive correlation between 5hmC and transcription was also observed at genes with gene body 5hmC peaks, however this effect was less pronounced, likely due to challenges in associating distal 5hmC peaks with their target genes **(Fig. S4E)**. In sum, these results provide proof for the presence of 5hmC at regulatory regions of developmental genes in invertebrate deuterostome genomes and suggest that active removal of 5mC through the 5hmC intermediate might be crucial for their activation.

**Figure 4.**
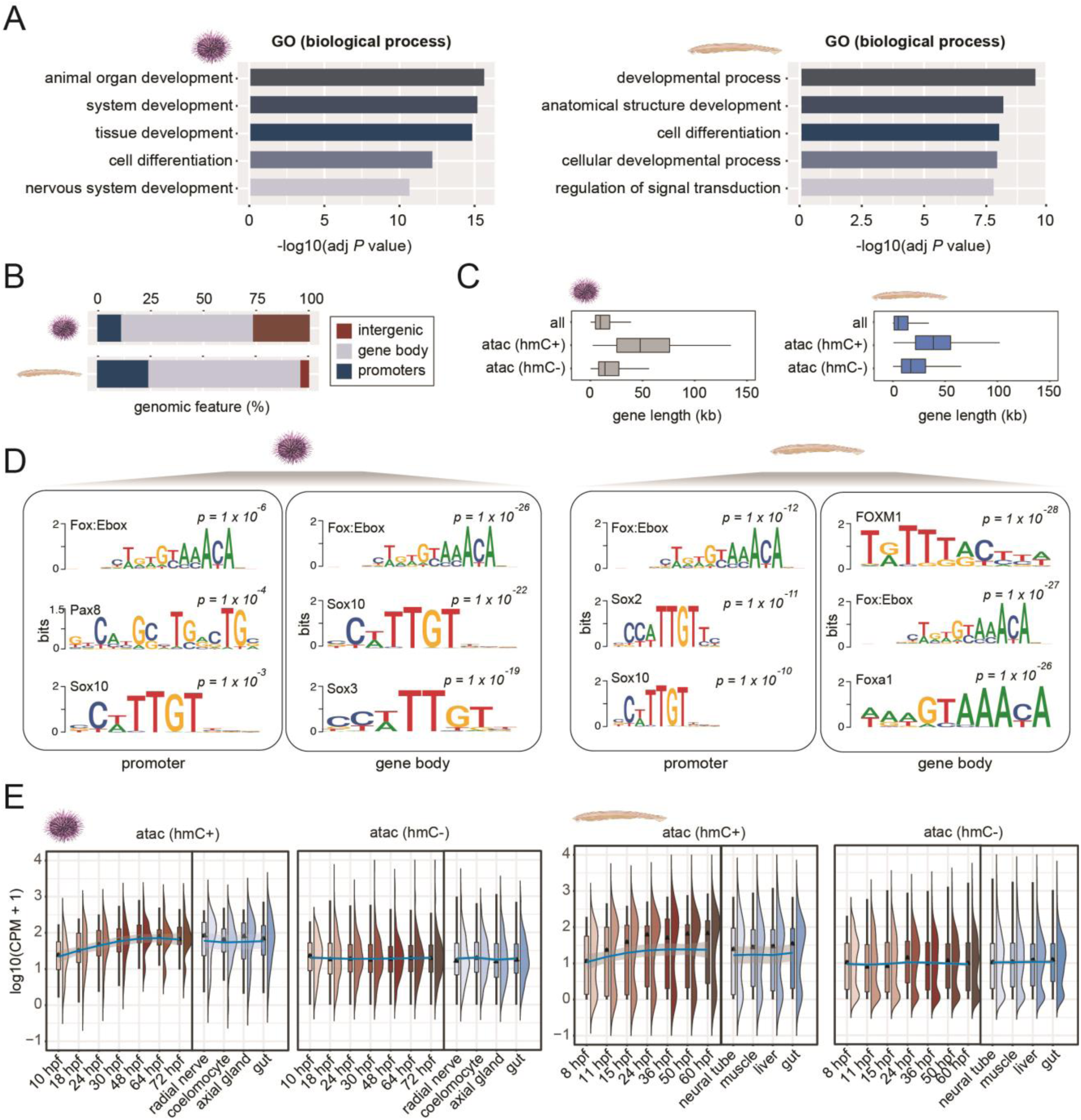
Developmental expression of 5hmC-marked genes. **(A)** Gene ontology analysis of genes harbouring 5hmC-marked ATAC-seq peaks at their promoters or gene bodies in sea urchin and lancelet. **(B)** Percentage of 5hmC-marked ATAC-seq peaks at different genomic regions. **(C)** Length of the genes harbouring promoter-associated 5hmC-marked ATAC-seq peaks (promoter atac hmC+) and non-5hmC ATAC-seq peaks (promoter atac hmC-). **(D)** HOMER motif enrichment analysis of atac hmC+ regions associated with either gene promoters or gene bodies. **(E)** Expression dynamics of promoter atac hmC+ genes as compared to promoter atac hmC-genes in sea urchin and lancelet embryos and adult tissues.

### Conserved regulatory logic of 5-hydroxymethylcytosine-marked genes

Previously, we have described widespread TET-mediated demethylation of regulatory regions during the vertebrate phylotypic period, which can be defined as the most conserved stage of vertebrate embryogenesis^19^. These phylotypic DMRs (phylo-DMRs), coincided with developmental DNA demethylation and were marked by 5hmC in zebrafish, *Xenopus* and mouse embryos. To explore the extent to which genes regulated by 5hmC in invertebrate deuterostomes coincide with genes linked to vertebrate 5hmC, we first identified zebrafish orthologues of 5hmC-marked genes in sea urchin and lancelet, defined their regulatory landscapes^60^, selected the zebrafish orthologs enriched in 5hmC, and assessed their overlap **(Fig. S5A)**. This analysis resulted in 148, and 343 zebrafish 5hmC-marked genes, which overlapped lancelet and sea urchin 5hmC-marked genes, respectively. Having identified this notable agreement in 5hmC regulation, we next wanted to explore in more detail the regulatory logic behind 5hmC gene marking. To that end we selected zebrafish orthologues of sea urchin 5hmC-marked genes and stratified ATAC-seq-associated regulatory elements in the zebrafish genome within each gene’s landscape^60^ into three categories depending on phylo-DMR presence and 5hmC enrichment **(Fig. S5B)**: group 1 (ATAC-seq peaks overlapping a phylo-DMR and displaying 5hmC enrichment), group 2 (ATAC-seq peaks displaying 5hmC enrichment without phylo-DMR overlap), and group 3 (ATAC-seq peaks without 5hmC enrichment or phylo-DMR overlap). We next explored the transcriptional profiles of zebrafish orthologs linked to these three groups and found that phylo-DMR-linked genes (group 1) were characterised by peaks of expression surrounding the phylotypic period (24hpf), whereas 5hmC-marked genes without phylo-DMR association (group 2) were developmentally upregulated at later embryonic stages (48-72hpf) and exhibited strongest expression in the adult brain (**Fig. 5A**). Genes that were not enriched in 5hmC or phylo-DMRs (group 3) did not display such notable developmental dynamics (**Fig. 5A**). To obtain further insight into the spatio-temporal regulation of these orthologs, we analysed single cell resolution transcriptomes of the zebrafish phylotypic stage embryo (24hpf) (**Fig. 5B**)^66^. We found that group 1 genes were expressed broadly in cell-types originating from diverse embryonic layers, whereas group 2 genes were expressed predominantly in neurons, in line with their bulk RNA-seq expression profile (**Fig. 5A-C, Fig. S5C**). Group 3 orthologues were characterised by a minor enrichment in the erythroid lineage. Given the strong link between 5hmC and loss of 5mC in non-vertebrate (**Fig. 2**) and vertebrate^19^ adult tissues, we next sought to interrogate the 5mC state of these three groups of regulatory elements in adult zebrafish^67^ **(Fig. 5D**). We found that both 5hmC-enriched ATAC peaks with and without phylo-DMR association are predominantly hypomethylated in most adult tissues (**Fig. 5D)**. Unlike group 2 elements, which are exclusively hypomethylated in adults, group 1 elements were already hypomethylated during the phylotypic period. Finally, group 3 elements that displayed no 5hmC enrichment coincided with peaks of broad and pervasive hypomethylation characteristic of CpG islands^68^. These data argue that most invertebrate deuterostome 5hmC-marked genes also display an association with this regulatory mark in vertebrates, which utilise 5hmC during organogenesis (group 1 elements) as well as during later stages of tissue differentiation (group 2 elements).

**Figure 5.**
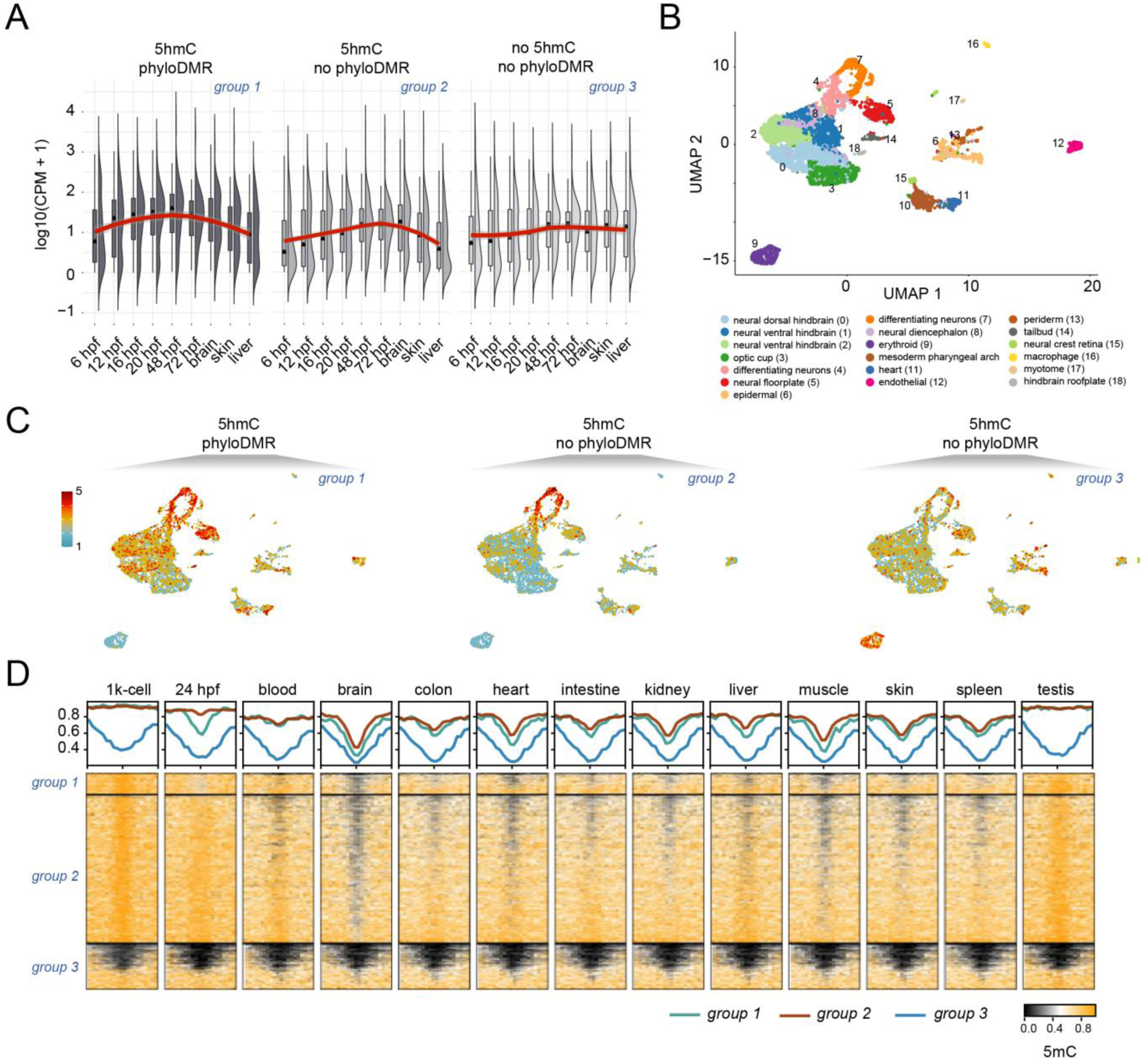
Conservation of developmental regulatory logic of 5hmC-marked genes. **(A)** Expression of 2R/3R-ohnologue zebrafish genes either associated with 5hmC-marked ATAC-seq peaks overlapping phylo-DMRs (5hmC/phylo-DMR), 5hmC-marked ATAC-seq peaks not overlapping phylo-DMRs (5hmC/no phylo-DMR) or non-5hmC ATAC-seq peaks not overlapping phylo-DMRs (no 5hmC/no phylo-DMR) in embryos and adult tissues. **(B)** Uniform Manifold Approximation and Projection (UMAP) projection of 7,738 individual cells from the 24hpf zebrafish embryo. Cells are colour-coded by tissue types. **(C)** z-transformed module scores of three gene sets: 5hmC/phylo-DMR, 5hmC/no phylo-DMR and no 5hmC/no phylo-DMR in 24hpf zebrafish embryo. Module scores depict the difference between the average gene expression of each gene set and random control genes. **(D)** Heatmaps showing DNA methylation profiles of 5hmC/phyloDMR, 5hmC/no phyloDMR and no 5hmC/no phyloDMR ATAC-seq peaks in 1K-cell and 24hpf zebrafish embryos as well as multiple adult tissues.

## Discussion

The goal of the current study was to explore the evolutionary conservation of TET-mediated DNA demethylation in non-vertebrate lineages by generating base-resolution profiles of developmental 5-hydroxymethylcytosine landscape dynamics. Firstly, we have shown that the expression of sea urchin and lancelet TET orthologues by and large resembles anamniote TET developmental expression profiles, which are characterised by peaks of TET activity during mid-development (phylotypic period)^19^. While TET proteins play important roles in mammalian pluripotency and reprogramming *in vitro*^21,64,65,69^, mouse triple TET knockouts developed normally until gastrulation^18^ indicative of reduced requirements for TET activity during *in vivo* pluripotency. Similarly, zebrafish homozygous TET knockouts generated by TALE technology^22^, or CRISPR/Cas9 F0 mutants^20^ and morphants^19^, all displayed developmental defects only during later embryonic stages. Our results thus suggest that active DNA demethylation removal and likely 5mC in general, are not required for the establishment of deuterostome pluripotency.

While the members of TET/JBP (J-protein binding) family have been identified across all domains of life, ranging from viruses to humans^33^, fairly little is known regarding the functions of these proteins in DNA demethylation beyond vertebrates^28,35,37^. For example, in *Drosophila*, the single identified TET orthologue has only been associated with RNA hydroxymethylation^34^ whereas detailed, base-resolution invertebrate 5-hydroxymethylcytosine maps have thus far only been generated for the demosponge *A. queenslandica* and the sea anemone *Nematostella vectensis*^31^. Interestingly however, in *A. queenslandica*, 5hmC was detectable at identical positions during embryonic and larval stages without displaying any signs of developmental loss^28^. This suggests that *A. queenslandica* either uses 5hmC as a stable gene-regulatory mark or that TET-mediated DNA demethylation occurs during later developmental transitions. In this work we conclusively demonstrate that genomic sites of 5hmC enrichment coincide with developmental 5mC loss and the presence of accessible chromatin, in line with the proposed roles of TET proteins and 5hmC in gene regulation^70^. Specifically, we detect 5hmC at both proximal and distal regulatory regions and show that 5hmC presence and developmental loss of 5mC coincide with activation of developmental genes. We have previously demonstrated that active DNA demethylation of enhancers associated with key developmental pathways is a hallmark of vertebrate development^19^. Here we further expand on these observations to conclude that TET activity and active DNA demethylation of regulatory regions during mid-developmental stages could form a conserved developmental signature of diverse animal phyla. It is worth noting however, that the current study is limited to invertebrate deuterostomes from the phylum Echinodermata (*S. purpuratus*) and Chordata (*B. lanceolatum*), making it thus challenging to extrapolate our findings beyond these two lineages. Deeply conserved DNA demethylation toolkits consisting of TET proteins and DNA repair machinery as well as canonical gene body 5mC patterns observed in most animals^28,33^ would support widespread usage of TETs and 5hmC for gene regulation. Nevertheless, transcriptome profiling of multiple developmental stages across ten different phyla demonstrated divergence in the usage of signalling pathways and transcription factors during mid-development^71^. It remains to be seen how prevalent TET-mediated regulation of developmental genes is in other animal lineages, and if 5mC/5hmC regulatory logic could be ancestral to bilaterian animals. Finally, in the current study we have shown that 5hmC-regulated genes identified in invertebrate deuterosteomes also display 5mC and 5hmC-mediated regulation in the zebrafish genome and that most of such 5hmC-enriched genes are either linked to actively demethylated phylotypic enhancers or regulatory regions that become demethylated in diverse adult tissues, and in particular adult brain. Overall, our study provides novel insight into the 5mC/5hmC regulatory logic beyond the invertebrate-vertebrate boundary and demonstrates that active 5mC removal from regulatory regions during animal development predates vertebrate origins.

## Methods

### Genomic DNA extraction from sea urchin embryos and adult tissues

*Strongylocentrotus purpuratus* adult animals were collected in the San Diego Bay area, San Diego County, California (USA) and kept in a closed tank system with circulating diluted Mediterranean Sea water at 36 ppt and 14°C at Stazione Zoologica Anton Dohrn. Adult tube feet were collected by surgically removing them from the adult sea urchin attached to a glass beaker with sea water. To obtain embryos and larvae, sea urchin gametes were collected by vigorously shaking the adults. Eggs were collected by placing the sea urchin aboral side up onto a beaker filled with seawater. Sperm was collected with a p200 micropipette into a 1.5 mL tube, which was placed on ice. The eggs were filtered once through a 100 μm mesh filter to remove debris. 5 μL of sperm was diluted in 13 mL of seawater, then 20 drops of this dilution were added to the eggs to fertilize them. Fertilized eggs were incubated in seawater at 15°C until the desired stage. 25-50 μL of sea urchin larvae or embryos were collected for DNA extraction by using a 40 μm mesh filter and placed into 1.5 mL tubes. Tubes with adult tube feet, embryos and larvae were centrifuged at 500 g and the supernatant was discarded. The resulting pellets were frozen by placing the tubes in liquid nitrogen and then stored at -80°C until further processing. The DNA extraction procedure was identical for sea urchin adult tube feet, embryos, and larvae. The frozen pellet was washed 3 times with 1X SSC in milliQ H2O by centrifuging at 500 x g for 5 minutes and removing the supernatant. The washed pellet was resuspended in 750 mL of lysis buffer (10 mM Tris pH 7.5, 1 mM EDTA, 1% SDS, 3.4% βmercaptoethanol, 660μg/mL Proteinase K) and incubated at 56°C for 2 hours at 300 rpm in a heat block. After cooling down, an equal amount of PCI (Phenol/Chlorophorm/Isoamylalcohol mixture in 25:24:1 proportion at neutral pH) was added and mixed by inverting the tube several times, followed by centrifugation at 16,000 x g for 10 minutes at 4°C to separate the phases. The top phase was collected and mixed with an equal volume of CIA (Chloroform/Isoamyl Alcohol mixture in 24:1 proportion). The solution was mixed by inversion and centrifuged at 16,000 x g for 10 minutes at 4°C. The top phase was collected and combined with 0.1 volume of 3M NaAc (pH 5.0) and 2 volumes of icecold 100% EtOH. The solution was mixed by inverting the tube several times. Precipitated DNA was collected by a p20 micropipette and placed in a 1.5 mL tube with 200 μL of 70% EtOH. This procedure was repeated twice to wash the DNA. The precipitated DNA pellet was then placed into an empty clean 1.5mL tube and air-dried to remove EtOH. Dried DNA was resuspended in 200 to 500 μL of 1x TE buffer (pH 8.0) depending on the amount of DNA.

### Genomic DNA extraction from lancelet embryos and adult tissues

Adults of *Branchiostoma lanceolatum* were collected at the Racou beach near Argelès-sur-Mer, France (latitude 42° 32′ 53′′ N, longitude 3° 03′ 27′′ E) with a specific permission delivered by the Prefect of Region Provence Alpes Côte d’Azur. *Branchiostoma lanceolatum* is not a protected species. Gametes were obtained after heat stimulation as previously described^72^ and were used for *in vitro* fertilization in Petri dishes filled with filtered seawater. Embryos were collected at the desired stages and placed in 1.5 mL tubes (at least 500 embryos per tube). The seawater was discarded after 1 minute of centrifugation at 13,000 rpm. Adults were immobilized in MgCl_2_ 7% for 5 minutes, placed in seawater and sacrificed by decapitation. The hepatic caecum was dissected and placed in 1.5 mL tubes. The embryos and adult tissues were processed using the DNeasy Blood and Tissue kit (QIAGEN) following the instructions provided by the manufacturer, with an incubation in lysis buffer at 56°C for 5 hours and using 200μL of elution buffer.

### MethylC-seq library preparation

MethylC-seq library preparation was performed as described previously^40^. Briefly, 1000 ng of genomic DNA extracted from sea urchin and lancelet embryos and adult tissues was spiked with unmethylated λ phage DNA (Promega) and sonicated to ∼300 bp fragments using a M220 focused ultrasonicator (Covaris) with the following parameters: peak incident power, 50W; duty factor, 20%; cycles per burst, 200; treatment time, 75 sec. Sonicated DNA was then purified, end-repaired using End-It™ DNA End-Repair Kit (Lucigen) and A-tailed using Klenow Fragment (3’→5’ exo-) (New England Biolabs) followed by the ligation of NEXTFLEX^®^ Bisulfite-Seq adapters. Bisulfite conversion of adaptor-ligated DNA was performed using EZ DNA Methylation-Gold Kit (Zymo Research). Library amplification (13 PCR cycles) was performed with KAPA HiFi HotStart Uracil+ DNA polymerase (Kapa Biosystems). Library size was determined by the Agilent 4200 Tapestation system. The libraries were quantified using the KAPA library quantification kit (Roche), yielding ∼10-20 nM.

### ACE-seq library preparation

ACE-seq library preparation for base resolution 5-hydroxymethylcytosine profiling was performed as previously described^41,73^. Briefly, 100 ng of genomic DNA extracted from sea urchin, lancelet, and zebrafish embryos and adult tissues was spiked with 1 ng of CpG-methylated λ phage DNA (Wisegene) and 0.5 ng of all-C hydroxymethylated pUC19 plasmid DNA (Wisegene) and sonicated to ∼300 bp fragments using a M220 focused ultrasonicator (Covaris), with the following parameters: peak incident power, 50W; duty factor, 20%; cycles per burst, 200; treatment time, 75 sec. Sonicated DNA (125 μL total volume) was concentrated to 16.6 μL using AMPure beads (1 volume DNA : 2 volumes AMPure beads). 5hmC protection reaction was performed using β-glucosyltransferase (10U/μL, New England Biolabs) in Cutsmart Buffer according to manufacturer’s instructions at 37°C for 1 hr, followed by a 10 min incubation at 65°C. The DNA was denatured in DMSO at 95°C for 10 min and immediately placed on dry ice for 2 mins allowing the samples to freeze. C and 5mC deamination reaction was performed using the APOBEC3A enzyme (NEBNext® Enzymatic Methyl-seq Kit, New England Biolabs) with the following ramping conditions: 4°C for 10 min, 4°C - 50°C 2:15 min per degree of the ramp, 50°C for 10 min. Deaminated DNA was then purified using AMPure beads (1 volume DNA: 1 volume AMPure beads) and subjected to low input library preparation using Accel-NGS Methyl-Seq DNA kit (Swift Biosciences). Briefly, DNA was denatured and subjected to an adaptase reaction, followed by primer extension, adapter ligation, and 15 cycles of indexing PCR. Library size and consistency was determined by the Agilent 4200 Tapestation system. The libraries were quantified using the KAPA library quantification kit (Roche).

### hMeDIP-seq library preparation

The hMeDIP assay was performed according to the manufacturer’s instructions (Active Motif, hMeDIP, Cat No 55010) on sea urchin 72hpf, lancelet 36hpf, and zebrafish 24hpf embryos. DNA was sonicated with the Covaris sonicator to generate ∼300 bp fragments. Briefly, 200-400 ng of fragmented DNA was spiked with either unmethylated, 5mC methylated or 5hmC hydroxymethylated 338-bp PCR product of APC genomic locus. The DNA was denatured for 10 min at 95°C and immunoprecipitated overnight at 4°C with 4 μL of 5hmC monoclonal antibody (Active Motif Cat No 55010). To allow selective enrichment of immune-captured DNA fragments, the mixture was incubated with 25 μL of Protein G magnetic beads for 2 h at 4°C prior to washing of unbound DNA fragments. Bound hydroxymethylated DNA was eluted, treated with proteinase K and purified by Phenol-chloroform/isoamyl alcohol extraction and ethanol precipitation. hmC-containing DNA was subjected to library preparation using the TruSeq ChIP Sample Preparation kit, (Illumina). The specificity of the hMeDIP assay was validated by qPCR of the unmethylated, methylated and hydroxymethylated spike-in APC controls.

### ATAC-seq library preparation

ATAC-seq assays were performed on sea urchin embryos as previously described^74,75^. For each of the two ATAC-seq replicates, approximately 45,000 cells (corresponding to ∼135 embryos at 48 hpf) were lysed in cold lysis buffer (10 mM Tris pH 7.4, 10 mM NaCl, 3 mM MgCl2 and 0.2% NP40/Igepal) after removing the seawater and washing the embryos with filtered ice-cold artificial seawater three times by centrifuging at 500 x g. The resultant nuclei were then incubated for 30 min at 37°C with an in-house made Tn5 enzyme and purified with the Qiagen MinElute kit (Qiagen, 28004). PCR reactions for each replicate were performed with 15 cycles using Ad1F and Ad2.2R primers and NEBNext High-Fidelity 2X PCR Master Mix (New England Labs Cat #M0541). The resulting libraries were multiplexed and sequenced on the HiSeq 4000 sequencer.

### MethylC-seq data analysis

Sea urchin and lancelet MethylC-seq libraries were sequenced on the Illumina HiSeqX platform (150 bp, PE). Sequenced reads in FASTQ format were trimmed using Trimmomatic (ILLUMINA-CLIP:adapter.fa:2:30:10 SLIDINGWINDOW:5:20 LEADING:3 TRAILING:3 MINLEN:50). Trimmed reads were mapped to Spur_v3.1, Bl71nemr and danRer10 genome references for sea urchin, lancelet, and zebrafish, respectively, (all containing the λ phage genome as chrLambda) using WALT^76^ with the following settings: -m 10 -t 24 -N 10000000 -L 2000. Mapped reads in SAM format were converted to BAM and indexed using SAMtools^77^. Optical and PCR duplicates were removed using Picard Tools function *MarkDuplicates REMOVE_DUPLI-CATES=true*. Methylated and unmethylated cytosines at each genomic CpG position were called using MethylDackel v0.3.0. Additional *MethylDackel extract* parameters *--minOppositeDepth 5 --maxVariantFrac 0*.*5* were used for the genotype correction (in order to exclude C-to-T nucleotide transitions). Additionally, sequencing read boundaries obtained from *MethylDackel mbias* were reviewed and fed into *MethylDackel extract --OT a,b,c,d, --OB a,b,c,d* parameters. For MethylC-seq, bisulfite conversion efficiency was estimated from the λ phage spike-in control.

### ACE-seq data analysis

Sea urchin, lancelet, zebrafish and mouse ACE-seq libraries were sequenced on Illumina HiSeqX and NovaSeq 6000 platforms (150 bp, PE). Adapter clipping and read quality trimming was performed using Trimmomatic (ILLUMINACLIP-:NEXTflex.fa:2:30:10 SLIDINGWINDOW:5:20 LEADING:5 TRAILING:5 MINLEN:50 CROP:140 HEADCROP:10)^78^. These settings allow for the removal of low complexity Adaptase tails. Trimmed reads were mapped to Spur_v3.1, Bl71nemr, danRer10 and mm10 genome references for sea urchin, lancelet, zebrafish and mouse^41^, respectively, (all containing the λ phage genome as chrLambda and pUC19 plasmid genome as chrPUC) using WALT^76^ with the following settings: -m 10 -t 24 -N 10000000 -L 2000. Mapped reads in SAM format were converted to BAM and indexed using SAMtools^77^. Optical and PCR duplicates were removed using Picard Tools’s function *MarkDuplicates REMOVE_DUPLICATES=true*. Reads with more than three consecutive non-converted cytosines in the CC, CA and CT context were removed using Picard Tools’ function *FilterSamReads*. In order to determine sequencing read boundaries displaying uniform hydroxymethylation rate and to increase the accuracy of CpG hydroxymethylation calling, we ran the methylation bias command: *MethylDackel mbias*. The numbers of hydroxymethylated and unmethylated cytosines at each genomic CpG position were called using the *MethylDackel extract genome*.*fasta input*.*bam -o output --mergeContext*. Additional *MethylDackel extract* parameters *--minOppositeDepth 5 --maxVariantFrac 0*.*5* were used for genotype correction. Additionally, sequencing read boundaries obtained from *MethylDackel mbias* were reviewed and fed into *MethylDackel extract --OT a,b,c,d, --OB a,b,c,d* parameters.

### Statistical analysis of hydroxymethylation calling

Statistical analysis of the accuracy of ACE-seq 5hmC calling at a given CpG site was performed following the assumption that the number of “C” reads (N_c_) for each CpG site follows a binomial distribution *N*_*c*_ *∼ binomial(p, n)* where *p* (probability distribution) is the 5mC-to-T non-conversion rate and *n* is sequencing depth (number of “C” + “T” reads per each CpG site). Proportion test (R *prop*.*test*) was utilized to test the null hypothesis H0 that the observed proportion (the number of “C” reads at a given CpG site) equals the expected proportion (p) and the alternative hypothesis H1 that that the observed proportion is greater than the expected proportion. The following parameters were used: *prop*.*test(x, n, p = nonconv_rate, alternative = “greater”)*, where *x* is a vector of counts of successes (number of “C” reads for each CpG site), *n* - a vector of counts of trials (number of “C” + “T” reads per each CpG site, i.e coverage), *p* - a vector of probabilities of success (5mC-to-T non-conversion rate). Non-conversion rate of each ACE-seq library was estimated using the M.SssI CpG methyltransferase 5mCG-methylated λ phage spike-in controls. The non-conversion rate is the proportion of “C” reads at methylated CpG sites of the 5mCG λ phage spike-in control being incorrectly retained as cytosines due to inefficient APOBEC3A-mediated deamination of 5mC. The statistical analysis was restricted to CpG sites covered by at least 10 reads.

### Identification of differentially methylated regions (DMRs)

Differentially methylated CpG sites between consecutive embryonic and adult stages were identified using the *DMLtest* function of the DSS package^79^. Differentially methylated CpG sites located within 50 bp from each other were joined into regions (DMRs), which were filtered to harbour at least 5 differentially methylated CpGs that span at least 50 bp. DMRs containing more than 25% of CpGs within them with sequencing coverage less than 5x were excluded from the analysis. The remaining DMRs were defined as hypomethylated or hypermethylated when both replicates displayed at least 20% average methylation difference between consecutive developmental stages.

### hMeDIP-seq data analysis

hMeDIP-seq libraries from sea urchin 72hpf, lancelet 36hpf, and zebrafish 24hpf embryos were sequenced on Illumina HiSeq X platform. hMeDIP-seq raw data was processed using the NGSANE framework v0.5.2.0^80^. Briefly, sequenced hMeDIP-seq reads in FASTQ format were trimmed using Trimmomatic (ILLUMINACLIP:TruSeq3-PE.fa:2:30:10 SLIDINGWINDOW:5:20). Trimmed reads were mapped to Spur_v3.1, Bl71nemr and danRer10 reference genomes for sea urchin, lancelet and zebrafish, respectively, using bowtie2 v2.1.0 with the following parameters: *bowtie2 --end-to-end -R 2 -p 10 -N 1 --very-sensitive -X 1000 --no-mixed - -no-discordant*^81^. Optical and PCR duplicates were removed using Picard Tools function *MarkDuplicates REMOVE_DUPLICATES=true*. hMeDIP-seq signal enrichment over ATAC-seq peaks was plotted using deepTools2 *computeMatrix reference-point* and *plotHeatmap* functions.

### ATAC-seq data analysis

Sea urchin ATAC-seq data was generated as part of this study, whereas lancelet and zebrafish ATAC-seq data were downloaded from GSE106428. Sequenced ATAC-seq reads in FASTQ format were trimmed using Trimmomatic software (ILLUMINACLIP:TruSeq3-PE.fa:2:30:10 SLIDINGWINDOW:5:20 LEADING:3 TRAILING:3 MINLEN:25). Properly paired trimmed read pairs were mapped to Spur_v3.1, Bl71nemr and danRer10 genome references for sea urchin, lancelet, and zebrafish, respectively, using bowtie2 v2.1.0 with the following parameters: *bowtie2 -p 10 -N 1 --very-sensitive -X 2000 --no-mixed --no-discordant*^81^. Mapped reads in SAM format were converted to BAM and indexed using SAMtools^77^. Optical and PCR duplicates were removed using Picard Tools function *MarkDuplicates REMOVE_DUPLICATES=true*. ATAC-seq fragment length distribution was plotted using deepTools2^82^ *bamPEFragmentSize --histogra*. Removal of mono-, di- and trinucleosome-bound DNA fragments (and retention of the nucleosome-free fragments) and read adjustment +4 bp -5 bp for positive and negative strand, respectively, was performed using deepTools2 *alignmentSieve -- ATACshift --maxFragmentLength 100*. Peaks were called using MACS2^83^ with the following parameters: *macs2 callpeak -t $input_bam -f BAMPE -g ${genome_size} --nomodel --nolambda - -outdir ${OUTPUT_DIR} -n $name --call-summits - B -q 0*.*05*.

### RNA-seq data analysis

Sea urchin RNA-seq data was downloaded from PRJNA81157, whereas lancelet and zebrafish RNA-seq data were downloaded from GSE106430 / PRJNA416866. Illumina adapter trimming was performed using TrimGalore v0.4.0 (https://github.com/FelixKrueger/TrimGalore). Trimmed sequence reads were aligned to the Spur_v3.1, Bl71nemr and danRer10 reference genomes for sea urchin, lancelet and zebrafish, respectively, using STAR v2.4.0d^84^. Quantification of transcript abundance was performed using RSEM v1.2.2^85^. TET genes analysed in this study correspond to the following IDs: WHL22.387614 (SPU_027517) for sea urchin, BL00006 for the European lancelet, and ENSDARG00000075230,ENSDARG00000076928, ENSDARG00000062646 for zebrafish.

### TET protein sequence and structure conservation analysis

Sea urchin TET protein sequence was retrieved from The Universal Protein Resource (UniProt): UniProtKB - A0A7M7NF23 Methylcytosine dioxygenase TET (A0A7M7NF23_STRPU). Florida lancelet TET protein sequence was retrieved from https://ftp.ncbi.nlm.nih.gov/genomes/all/annotation_releases/7739/100/GCF_000003815.2_Bfl_VNyyK/GCF_000003815.2_Bfl_VNyyK_protein.faa.gz, XP_035668917.1 methylcytosine dioxygenase TET1-like isoform X1 [Branchiostoma floridae]. Fruit fly, zebrafish, mouse, and human TET protein sequences were retrieved from UniProt. To predict TET protein domain structure, amino acid sequences of target sTET, bTET and zebrafish TET1, TET2, TET3 proteins in FASTA format were used as input for HMMER (http://hmmer.org/) and SWISS-MODEL^86^. Multiple alignment of TET proteins amino acid sequences was performed using UniProt. The MSA comparison was visualised using Jalview 2.11.1.4^87^. SWISS-MODEL was used to perform protein structure homology modelling^86^. First, amino acid sequences of target sTET, bTET and zebrafish TET1, TET2, TET3 proteins in FASTA format were used as input. 5d9y.1.A (crystal structure of human TET2-5fC complex) and 5exh.1.C (crystal structure of mTET3-CxxC domain in complex with 5carboxylcytosine DNA) were identified as the top ranked templates based on GMQE (Global Model Quality Estimate), which provides a quality estimate combining the properties from the target-template alignment and the template structure. Sequence identity between the input and 5d9y.1.A template was 57% and 64% for sea urchin sTET and lancelet sTET, respectively; and 71%, 75%, 67% for zebrafish TET1, TET2, and TET3, respectively. 5d9y.1.A template was used to build a 3D model of the TET N-terminal catalytic domain. Model quality was assessed by QMEANDisCo global (the entire structure) quality score, which is a quality measurement ranging between 0 to 1; the closer the score to 1, the higher the expected quality. QMEANDisCo global score was 0.72 and 0.70 for sea urchin sTET and lancelet sTET, respectively; and 0.62, 0.57 and 0.80 for zebrafish TET1, TET2, and TET3, respectively.

### Analysis of developmentally hydroxymethylated zebrafish gene orthologs

To assess the temporal and spatial expression of zebrafish orthologs of sea urchin genes linked to 5hmC enrichment (ATAC hmC+ genes), we first identified orthologous genes by converting sea urchin and zebrafish gene IDs to *Drosophila melanogaster* gene IDs using ENSEMBL BioMART tool. Next, sea urchin genes harbouring ATAC hmC+ peaks within their promoters or gene bodies were identified. Zebrafish genes, orthologous to sea urchin ATAC hmC+ genes, were used in the downstream analysis. To compare developmental expression dynamics of zebrafish genes possessing distinct regulatory landscapes after three rounds of whole genome duplication, we identified zebrafish 2R/3R-ohnologues among zebrafish genes, orthologous to the sea urchin ATAC hmC+ genes. 2R and 3R ohnologues (Intermediate: *q*-score (outgroups) < 0.01 AND *q-*score (self-comparison) < 0.01) were downloaded from http://ohnologs.curie.fr^88^. Ohnologues were first split into three groups: (i) genes associated with 5hmC-marked ATAC-seq peaks overlapping phylo-DMRs (5hmC/phylo-DMR), (ii) 5hmC-marked ATAC-seq peaks not overlapping phylo-DMRs (5hmC/no phylo-DMR) and (iii) non-5hmC ATAC-seq peaks not overlapping phylo-DMRs (no 5hmC/no phylo-DMR) in their distal regulatory domains. Zebrafish distal regulatory domains were defined using GREAT gene regulatory domain definition^60^. Briefly, basal regulatory domains were defined as 5 kb upstream and 1 kb downstream of the TSS, regardless of the neighbouring genes. To define distal regulatory domains, basal regulatory domains of each gene were extended upstream and downstream to the neighbouring gene’s basal domain but no more than 1000 kb in each direction. Phylo-DMRs were downloaded from^19^. 24hpf and 48hpf ATAC-seq peaks were downloaded from DANIO-CODE (ATAC-seq_Skarmeta_Lab_0001AS, https://danio-code.zfin.org)^19,89^. Only the ohnolog-ues with at least two different regulatory landscapes (5hmC/phylo-DMR, 5hmC/no phylo-DMR or no 5hmC/no phylo-DMR) were considered in the downstream bulk RNA-seq and scRNA-seq analysis.

### scRNA-seq data analysis

Zebrafish 24hpf embryo single cell RNA-seq InDrop data was downloaded from GSE112294^66^. The data was converted to a file format that resembles 10X Chromium CellRanger output using the indRop R package (https://github.com/caleblareau/indRop) and processed using the standard *Seurat* pipeline with the parameters suggested by developers^90^. Briefly, single cells with less than 200 detected genes, more than 4000 detected genes, indicative of empty droplets and duplicate cells in the droplet, respectively, were discarded. Cells with total UMI (transcript) counts less than 1000 and more than 15,000 were excluded from the analysis. Read counts were normalized by a factor of 10,000 and log transformed. Highly variable features were identified using *FindVariableFeatures* function. The number of UMIs per cell was regressed out using Seurat’s *ScaleData* function. *RunPCA* was used to perform linear dimensionality reduction on scaled data using determined highly variable features. Cell clustering was performed using *FindNeighbors(object, reduction = “pca”, dims = 1:20)* and *FindClusters(object, resolution = 0*.*5, algorithm = 1)*. Non-linear dimensionality reduction was performed using Uniform Manifold Approximation and Projection: *RunUMAP(object, reduction = “pca”, dims = 1:20)*.

### scRNA-seq downstream analysis

*FindAllMarkers* function was used to define gene markers for every cluster in the 24hpf zebrafish embryo. Top 20 genes identified as cluster features^66^ were matched with the obtained gene markers and used to assign cell clusters to different tissue types. *AddModuleScore* function was used to define module scores for a set of genes harbouring a 5hmC peak and a phyloDMR (5hmC / phyloDMR), and genes harbouring a 5hmC peak not overlapping a phyloDMR (5hmC / no phyloDMR), and genes without a 5hmC peak and without a phyloDMR (no 5hmC / no phyloDMR). Positive module score indicates that a given module of genes is more highly expressed in a certain cell than what would be expected from the average expression of this gene module across all cells in the population. Module score was then z-transformed and used in the *FeaturePlot*. To generate *‘upset’* plots, *PrctCellExpringGene* function was used to calculate the percentage of cells expressing each of the genes in the 5hmC/phyloDMR, 5hmC/no phyloDMR and no 5hmC/no phyloDMR gene sets. Genes with non-zero expression counts in more than 25% of cells in any given cluster, were deemed expressed in that cluster. *UpSetR* package upset function was used to display the number of genes in the gene set expressed per cluster, as well as the number of genes expressed across different numbers of clusters.

### DNA sequence motif analysis

Motif analysis of the 5hmC-marked ATAC-seq regions (+/-250 bp from the ATAC-seq peak summit) associated with promoters or gene bodies was performed using HOMER *findMotifsGenome*.*pl* using the default HOMER background normalised for GC content^91^. Motifs were visualised using R package *ggseqlogo*^92^.

## Data availability

MethylC-Seq, ACE-seq and ATAC-seq data generated in this study are available from NCBI Gene Expression Omnibus^93^ (GEO SuperSeries accession number GSE188334). Specifically, sea urchin and lancelet MethylC-seq are available under GSE188333. Sea Urchin, lancelet and zebrafish ACE-seq data are available under GSE188331. Sea Urchin, lancelet and zebrafish hMeDIP-seq data are available under GSE188332. Sea urchin ATAC-seq data are available under GSE186363. Public lancelet and zebrafish ATAC-seq data are available under GSE106428. Public sea urchin RNA-seq data are available under PRJNA81157. Public lancelet and zebrafish RNA-seq data are available under GSE106430 / PRJNA416866. Public single cell RNA-seq data of zebrafish 24 hpf embryo is available under GSE112294.

## Acknowledgements

The authors would like to thank Alex de Mendoza for critical reading of the manuscript, Patrick Leahy for providing adult sea urchins, and Davide Caramiello for sea urchin care. Australian Research Council (ARC) Discovery Project [DP190103852] to OB supported this work. Drawings of adult sea urchin, lancelet, and zebrafish were created with BioRender.com.

## Author contributions

OB and KS designed the study. MIA, HE and JLGS contributed to concept and study design. Sea urchin embryo and adult tissue collection and DNA extraction were performed by DV and MIA. Lancelet embryo and adult tissue collection and DNA extraction were performed by SB and HE. MethylC-seq library preparation and sequencing was carried out by PED and SER. ACE-seq library preparation and sequencing was carried out by KS. Sea urchin ATAC-seq library preparation was carried out by MSM, DV, MIA and IM and sequencing was carried out by MSM and IM. KS performed data analysis. OB and RJW participated in data analysis. KS and OB wrote the manuscript. All authors discussed the results and commented on the manuscript.

## References

1 Davidson, E. H. et al. A genomic regulatory network for development. Science 295, 1669–1678, doi:10.1126/science.1069883 (2002).

2 Woolfe, A. et al. Highly conserved non-coding sequences are associated with vertebrate development. PLoS Biol 3, e7, doi:10.1371/journal.pbio.0030007 (2005).

3 Bogdanovic, O. et al. Dynamics of enhancer chromatin signatures mark the transition from pluripotency to cell specification during embryogenesis. Genome Res 22, 2043–2053, doi:10.1101/gr.134833.111 (2012).

4 Schwaiger, M. et al. Evolutionary conservation of the eumetazoan gene regulatory landscape. Genome Res 24, 639–650, doi:10.1101/gr.162529.113 (2014).

5 Schubeler, D. Function and information content of DNA methylation. Nature 517, 321–326, doi:10.1038/nature14192 (2015).

6 Lee, H. J., Hore, T. A. & Reik, W. Reprogramming the methylome: erasing memory and creating diversity. Cell Stem Cell 14, 710–719, doi:10.1016/j.stem.2014.05.008 (2014).

7 Greenberg, M. V. C. Get Out and Stay Out: New Insights Into DNA Methylation Reprogramming in Mammals. Front Cell Dev Biol 8, 629068, doi:10.3389/fcell.2020.629068 (2020).

8 Rougier, N. et al. Chromosome methylation patterns during mammalian preimplantation development. Genes Dev 12, 2108–2113 (1998).

9 Ratnam, S. et al. Dynamics of Dnmt1 methyltransferase expression and intracellular localization during oogenesis and preimplantation development. Dev Biol 245, 304–314, doi:10.1006/dbio.2002.0628 (2002).

10 Cardoso, M. C. & Leonhardt, H. DNA methyltransferase is actively retained in the cytoplasm during early development. J Cell Biol 147, 25–32 (1999).

11 Mulholland, C. B. et al. Recent evolution of a TET-controlled and DPPA3/STELLA-driven pathway of passive DNA demethylation in mammals. Nat Commun 11, 5972, doi:10.1038/s41467-020-19603-1 (2020).

12 Nakamura, T. et al. PGC7/Stella protects against DNA demethylation in early embryogenesis. Nat Cell Biol 9, 64–71, doi:10.1038/ncb1519 (2007).

13 Wossidlo, M. et al. 5-Hydroxymethylcytosine in the mammalian zygote is linked with epigenetic reprogramming. Nat Commun 2, 241, doi:10.1038/ncomms1240 (2011).

14 Gu, T. P. et al. The role of Tet3 DNA dioxygenase in epigenetic reprogramming by oocytes. Nature 477, 606–610, doi:10.1038/nature10443 (2011).

15 Amouroux, R. et al. De novo DNA methylation drives 5hmC accumulation in mouse zygotes. Nat Cell Biol 18, 225–233, doi:10.1038/ncb3296 (2016).

16 Ito, S. et al. Tet proteins can convert 5-methylcytosine to 5-formylcytosine and 5-carboxylcytosine. Science 333, 1300–1303, doi:10.1126/science.1210597 (2011).

17 Kohli, R. M. & Zhang, Y. TET enzymes, TDG and the dynamics of DNA demethylation. Nature 502, 472–479, doi:10.1038/nature12750 (2013).

18 Dai, H. Q. et al. TET-mediated DNA demethylation controls gastrulation by regulating Lefty-Nodal signalling. Nature 538, 528–532, doi:10.1038/nature20095 (2016).

19 Bogdanovic, O. et al. Active DNA demethylation at enhancers during the vertebrate phylotypic period. Nat Genet 48, 417–426, doi:10.1038/ng.3522 (2016).

20 Ross, S. E. & Bogdanovic, O. Generation and Molecular Characterization of Transient tet1/2/3 Zebrafish Knockouts. Methods Mol Biol 2272, 281–318, doi:10.1007/978-1-0716-1294-1_17 (2021).

21 Hon, G. C. et al. 5mC oxidation by Tet2 modulates enhancer activity and timing of transcriptome reprogramming during differentiation. Mol Cell 56, 286–297, doi:10.1016/j.molcel.2014.08.026 (2014).

22 Li, C. et al. Overlapping Requirements for Tet2 and Tet3 in Normal Development and Hematopoietic Stem Cell Emergence. Cell Rep 12, 1133–1143, doi:10.1016/j.celrep.2015.07.025 (2015).

23 Koh, K. P. et al. Tet1 and Tet2 regulate 5-hydroxymethylcytosine production and cell lineage specification in mouse embryonic stem cells. Cell Stem Cell 8, 200–213, doi:10.1016/j.stem.2011.01.008 (2011).

24 Li, X. et al. Tet proteins influence the balance between neuroectodermal and mesodermal fate choice by inhibiting Wnt signaling. Proc Natl Acad Sci U S A 113, E8267–E8276, doi:10.1073/pnas.1617802113 (2016).

25 Blattler, A. et al. Global loss of DNA methylation uncovers intronic enhancers in genes showing expression changes. Genome Biol 15, 469, doi:10.1186/s13059-014-0469-0 (2014).

26 Suzuki, M. M. & Bird, A. DNA methylation landscapes: provocative insights from epigenomics. Nat Rev Genet 9, 465–476, doi:10.1038/nrg2341 (2008).

27 Bewick, A. J., Vogel, K. J., Moore, A. J. & Schmitz, R. J. Evolution of DNA Methylation across Insects. Mol Biol Evol 34, 654–665, doi:10.1093/molbev/msw264 (2017).

28 de Mendoza, A., Lister, R. & Bogdanovic, O. Evolution of DNA Methylome Diversity in Eukaryotes. J Mol Biol, doi:10.1016/j.jmb.2019.11.003 (2019).

29 Rae, P. M. & Steele, R. E. Absence of cytosine methylation at C-C-G-G and G-C-G-C sites in the rDNA coding regions and intervening sequences of Drosophila and the rDNA of other insects. Nucleic Acids Res 6, 2987–2995, doi:10.1093/nar/6.9.2987 (1979).

30 Simpson, V. J., Johnson, T. E. & Hammen, R. F. Caenorhabditis elegans DNA does not contain 5-methylcytosine at any time during development or aging. Nucleic Acids Res 14, 6711–6719 (1986).

31 de Mendoza, A. et al. Convergent evolution of a vertebrate-like methylome in a marine sponge. Nat Ecol Evol 3, 1464–1473, doi:10.1038/s41559-019-0983-2 (2019).

32 Neri, F. et al. Intragenic DNA methylation prevents spurious transcription initiation. Nature 543, 72–77, doi:10.1038/nature21373 (2017).

33 Parker, M. J., Weigele, P. R. & Saleh, L. Insights into the Biochemistry, Evolution, and Biotechnological Applications of the Ten-Eleven Translocation (TET) Enzymes. Biochemistry 58, 450–467, doi:10.1021/acs.biochem.8b01185 (2019).

34 Delatte, B. et al. RNA biochemistry. Transcriptome-wide distribution and function of RNA hydroxymethylcytosine. Science 351, 282–285, doi:10.1126/science.aac5253 (2016).

35 Wojciechowski, M. et al. Insights into DNA hydroxymethylation in the honeybee from in-depth analyses of TET dioxygenase. Open Biol 4, doi:10.1098/rsob.140110 (2014).

36 Cingolani, P. et al. Intronic non-CG DNA hydroxymethylation and alternative mRNA splicing in honey bees. BMC Genomics 14, 666, doi:10.1186/1471-2164-14-666 (2013).

37 Moroz, L. L. et al. The ctenophore genome and the evolutionary origins of neural systems. Nature 510, 109–114, doi:10.1038/nature13400 (2014).

38 Marletaz, F. et al. Amphioxus functional genomics and the origins of vertebrate gene regulation. Nature, doi:10.1038/s41586-018-0734-6 (2018).

39 Sea Urchin Genome Sequencing, C. et al. The genome of the sea urchin Strongylocentrotus purpuratus. Science 314, 941–952, doi:10.1126/science.1133609 (2006).

40 Urich, M. A., Nery, J. R., Lister, R., Schmitz, R. J. & Ecker, J. R. MethylC-seq library preparation for base-resolution whole-genome bisulfite sequencing. Nat Protoc 10, 475–483, doi:10.1038/nprot.2014.114 (2015).

41 Schutsky, E. K. et al. Nondestructive, base-resolution sequencing of 5-hydroxymethylcytosine using a DNA deaminase. Nat Biotechnol, doi:10.1038/nbt.4204 (2018).

42 Putnam, N. H. et al. The amphioxus genome and the evolution of the chordate karyotype. Nature 453, 1064–1071, doi:10.1038/nature06967 (2008).

43 Xu, Y. et al. Tet3 CXXC domain and dioxygenase activity cooperatively regulate key genes for Xenopus eye and neural development. Cell 151, 1200–1213, doi:10.1016/j.cell.2012.11.014 (2012).

44 Jessop, P., Ruzov, A. & Gering, M. Developmental Functions of the Dynamic DNA Methylome and Hydroxymethylome in the Mouse and Zebrafish: Similarities and Differences. Front Cell Dev Biol 6, 27, doi:10.3389/fcell.2018.00027 (2018).

45 Holland, P. W., Garcia-Fernandez, J., Williams, N. A. & Sidow, A. Gene duplications and the origins of vertebrate development. Dev Suppl, 125–133 (1994).

46 Rasmussen, K. D. & Helin, K. Role of TET enzymes in DNA methylation, development, and cancer. Genes Dev 30, 733–750, doi:10.1101/gad.276568.115 (2016).

47 Long, H. K., Blackledge, N. P. & Klose, R. J. ZF-CxxC domain-containing proteins, CpG islands and the chromatin connection. Biochem Soc Trans 41, 727–740, doi:10.1042/BST20130028 (2013).

48 Zhang, W. et al. Isoform Switch of TET1 Regulates DNA Demethylation and Mouse Development. Mol Cell 64, 1062–1073, doi:10.1016/j.molcel.2016.10.030 (2016).

49 Tu, Q., Cameron, R. A., Worley, K. C., Gibbs, R. A. & Davidson, E. H. Gene structure in the sea urchin Strongylocentrotus purpuratus based on transcriptome analysis. Genome Res 22, 2079–2087, doi:10.1101/gr.139170.112 (2012).

50 Bertrand, S. et al. The Ontology of the Amphioxus Anatomy and Life Cycle (AMPHX). Front Cell Dev Biol 9, 668025, doi:10.3389/fcell.2021.668025 (2021).

51 Carvalho, J. E. et al. An Updated Staging System for Cephalochordate Development: One Table Suits Them All. Front Cell Dev Biol 9, 668006, doi:10.3389/fcell.2021.668006 (2021).

52 Xu, X. et al. Evolutionary transition between invertebrates and vertebrates via methylation reprogramming in embryogenesis. Natl Sci Rev 6, 993–1003, doi:10.1093/nsr/nwz064 (2019).

53 Hon, G. C. et al. Epigenetic memory at embryonic enhancers identified in DNA methylation maps from adult mouse tissues. Nat Genet 45, 1198–1206, doi:10.1038/ng.2746 (2013).

54 Yu, M. et al. Base-resolution analysis of 5-hydroxymethylcytosine in the mammalian genome. Cell 149, 1368–1380, doi:10.1016/j.cell.2012.04.027 (2012).

55 Skvortsova, K. et al. Comprehensive evaluation of genome-wide 5-hydroxymethylcytosine profiling approaches in human DNA. Epigenetics Chromatin 10, 16, doi:10.1186/s13072-017-0123-7 (2017).

56 Kamstra, J. H., Loken, M., Alestrom, P. & Legler, J. Dynamics of DNA hydroxymethylation in zebrafish. Zebrafish 12, 230–237, doi:10.1089/zeb.2014.1033 (2015).

57 Seritrakul, P. & Gross, J. M. Tet-mediated DNA hydroxymethylation regulates retinal neurogenesis by modulating cell-extrinsic signaling pathways. PLoS Genet 13, e1006987, doi:10.1371/journal.pgen.1006987 (2017).

58 Skvortsova, K. & Bogdanovic, O. TAB-seq and ACE-seq Data Processing for Genome-Wide DNA hydroxymethylation Profiling. Methods Mol Biol 2272, 163–178, doi:10.1007/978-1-0716-1294-1_9 (2021).

59 Reimand, J., Kull, M., Peterson, H., Hansen, J. & Vilo, J. g:Profiler--a web-based toolset for functional profiling of gene lists from large-scale experiments. Nucleic Acids Res 35, W193–200, doi:10.1093/nar/gkm226 (2007).

60 McLean, C. Y. et al. GREAT improves functional interpretation of cis-regulatory regions. Nat Biotechnol 28, 495–501, doi:10.1038/nbt.1630 (2010).

61 Zeitlinger, J. & Stark, A. Developmental gene regulation in the era of genomics. Dev Biol 339, 230–239, doi:10.1016/j.ydbio.2009.12.039 (2010).

62 Vanzan, L. et al. High throughput screening identifies SOX2 as a super pioneer factor that inhibits DNA methylation maintenance at its binding sites. Nat Commun 12, 3337, doi:10.1038/s41467-021-23630-x (2021).

63 Yang, Y. A. et al. FOXA1 potentiates lineage-specific enhancer activation through modulating TET1 expression and function. Nucleic Acids Res 44, 8153–8164, doi:10.1093/nar/gkw498 (2016).

64 Ficz, G. et al. Dynamic regulation of 5-hydroxymethylcytosine in mouse ES cells and during differentiation. Nature 473, 398–402, doi:10.1038/nature10008 (2011).

65 Pastor, W. A. et al. Genome-wide mapping of 5-hydroxymethylcytosine in embryonic stem cells. Nature 473, 394–397, doi:10.1038/nature10102 (2011).

66 Wagner, D. E. et al. Single-cell mapping of gene expression landscapes and lineage in the zebrafish embryo. Science 360, 981–987, doi:10.1126/science.aar4362 (2018).

67 Yang, H. et al. A map of cis-regulatory elements and 3D genome structures in zebrafish. Nature 588, 337–343, doi:10.1038/s41586-020-2962-9 (2020).

68 Deaton, A. M. & Bird, A. CpG islands and the regulation of transcription. Genes Dev 25, 1010–1022, doi:10.1101/gad.2037511 (2011).

69 Hore, T. A. et al. Retinol and ascorbate drive erasure of epigenetic memory and enhance reprogramming to naive pluripotency by complementary mechanisms. Proc Natl Acad Sci U S A 113, 12202–12207, doi:10.1073/pnas.1608679113 (2016).

70 Pastor, W. A., Aravind, L. & Rao, A. TETonic shift: biological roles of TET proteins in DNA demethylation and transcription. Nat Rev Mol Cell Biol 14, 341–356, doi:10.1038/nrm3589 (2013).

71 Levin, M. et al. The mid-developmental transition and the evolution of animal body plans. Nature 531, 637–641, doi:10.1038/nature16994 (2016).

72 Fuentes, M. et al. Insights into spawning behavior and development of the European amphioxus (Branchiostoma lanceolatum). J Exp Zool B Mol Dev Evol 308, 484–493, doi:10.1002/jez.b.21179 (2007).

73 Wang, T. et al. Bisulfite-Free Sequencing of 5-Hydroxymethylcytosine with APOBEC-Coupled Epigenetic Sequencing (ACE-Seq). Methods Mol Biol 2198, 349–367, doi:10.1007/978-1-0716-0876-0_27 (2021).

74 Buenrostro, J. D., Giresi, P. G., Zaba, L. C., Chang, H. Y. & Greenleaf, W. J. Transposition of native chromatin for fast and sensitive epigenomic profiling of open chromatin, DNA-binding proteins and nucleosome position. Nat Methods 10, 1213–1218, doi:10.1038/nmeth.2688 (2013).

75 Magri, M. S. et al. ATAC-Seq for Assaying Chromatin Accessibility Protocol Using Echinoderm Embryos. Methods Mol Biol 2219, 253–265, doi:10.1007/978-1-0716-0974-3_16 (2021).

76 Chen, H., Smith, A. D. & Chen, T. WALT: fast and accurate read mapping for bisulfite sequencing. Bioinformatics 32, 3507–3509, doi:10.1093/bioinformatics/btw490 (2016).

77 Li, H. et al. The Sequence Alignment/Map format and SAMtools. Bioinformatics 25, 2078–2079, doi:10.1093/bioinformatics/btp352 (2009).

78 Bolger, A. M., Lohse, M. & Usadel, B. Trimmomatic: a flexible trimmer for Illumina sequence data. Bioinformatics 30, 2114–2120, doi:10.1093/bioinformatics/btu170 (2014).

79 Park, Y. & Wu, H. Differential methylation analysis for BS-seq data under general experimental design. Bioinformatics 32, 1446–1453, doi:10.1093/bioinformatics/btw026 (2016).

80 Buske, F. A., French, H. J., Smith, M. A., Clark, S. J. & Bauer, D. C. NGSANE: a lightweight production informatics framework for high-throughput data analysis. Bioinformatics 30, 1471–1472, doi:10.1093/bioinformatics/btu036 (2014).

81 Langmead, B. & Salzberg, S. L. Fast gapped-read alignment with Bowtie 2. Nat Methods 9, 357–359, doi:10.1038/nmeth.1923 (2012).

82 Ramirez, F. et al. deepTools2: a next generation web server for deep-sequencing data analysis. Nucleic Acids Res 44, W160–165, doi:10.1093/nar/gkw257 (2016).

83 Zhang, Y. et al. Model-based analysis of ChIP-Seq (MACS). Genome Biol 9, R137, doi:10.1186/gb-2008-9-9-r137 (2008).

84 Dobin, A. et al. STAR: ultrafast universal RNA-seq aligner. Bioinformatics 29, 15–21, doi:10.1093/bioinformatics/bts635 (2013).

85 Li, B. & Dewey, C. N. RSEM: accurate transcript quantification from RNA-Seq data with or without a reference genome. BMC Bioinformatics 12, 323, doi:10.1186/1471-2105-12-323 (2011).

86 Waterhouse, A. et al. SWISS-MODEL: homology modelling of protein structures and complexes. Nucleic Acids Res 46, W296–W303, doi:10.1093/nar/gky427 (2018).

87 Waterhouse, A. M., Procter, J. B., Martin, D. M., Clamp, M. & Barton, G. J. Jalview Version 2--a multiple sequence alignment editor and analysis workbench. Bioinformatics 25, 1189–1191, doi:10.1093/bioinformatics/btp033 (2009).

88 Singh, P. P. & Isambert, H. OHNOLOGS v2: a comprehensive resource for the genes retained from whole genome duplication in vertebrates. Nucleic Acids Res 48, D724–D730, doi:10.1093/nar/gkz909 (2020).

89 Baranasic, D. et al. (bioRxiv, 2021).

90 Satija, R., Farrell, J. A., Gennert, D., Schier, A. F. & Regev, A. Spatial reconstruction of single-cell gene expression data. Nat Biotechnol 33, 495–502, doi:10.1038/nbt.3192 (2015).

91 Heinz, S. et al. Simple combinations of lineage-determining transcription factors prime cis-regulatory elements required for macrophage and B cell identities. Mol Cell 38, 576–589, doi:10.1016/j.molcel.2010.05.004 (2010).

92 Wagih, O. ggseqlogo: a versatile R package for drawing sequence logos. Bioinformatics 33, 3645–3647, doi:10.1093/bioinformatics/btx469 (2017).

93 Edgar, R., Domrachev, M. & Lash, A. E. Gene Expression Omnibus: NCBI gene expression and hybridization array data repository. Nucleic Acids Res 30, 207–210 (2002).

