## Supplemental_Figures_1-5 for "Active DNA demethylation of developmental *cis*-regulatory regions predates vertebrate origins"

### Active DNA demethylation of developmental *cis*-regulatory regions predates the origin of vertebrates

#### Supplementary Figures 1-5

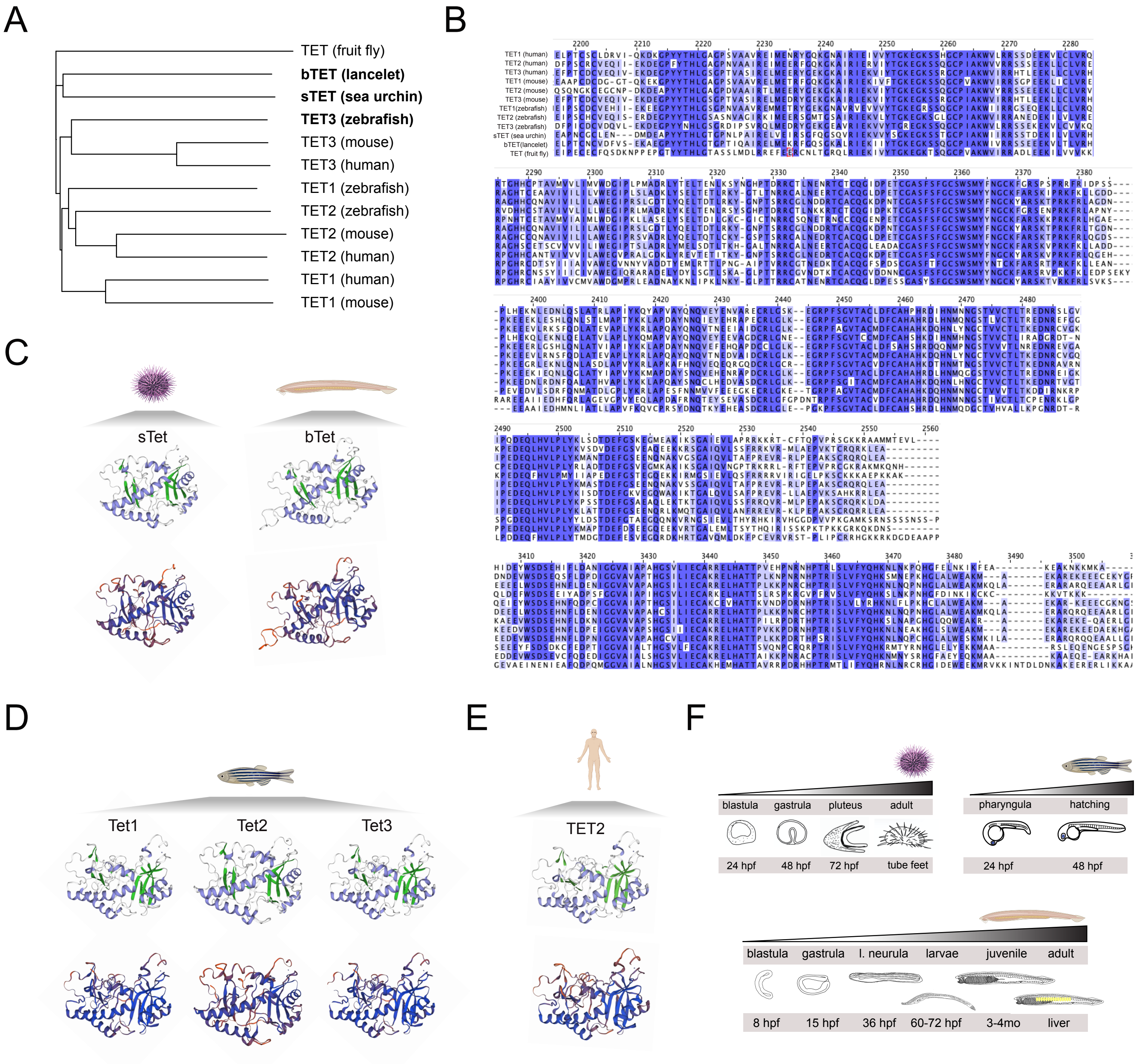


**Supplementary Figure 1. Conservation of TET protein sequence and catalytic domain structure. (A)** Cladogram depicting the phylogenetic relationships between fruit fly, sea urchin, lancelet, zebrafish, mouse, and human TET proteins. **(B)** Multiple sequence alignment of human, mouse and zebrafish TET1, TET2, TET3 as well as sea urchin, lancelet and fruit fly TET DSBH domains. DSBH domains harbour a large low-complexity insert, which is not shown. The colour of each amino acid indicates percentage identity (PID) with darker blue depicting higher PID and lighter blue - lower PID. **(C-E)** 3D models of methylcytosine dioxygenase domains of sea urchin sTet, lancelet bTet, zebrafish Tet1, Tet2, Tet3, and human TET2, performed using SWISS-MODEL. Top, 3D-model coloured by secondary structure: α-helix is depicted in blue and β-sheet in green. Bottom: 3D-model coloured by “Confidence”, QMEANDisCo local quality score depicting the expected similarity of each residue of the model to the native structure. High confidence is shown in blue, low confidence is shown in red. (**F**) Developmental stages of sea urchin, lancelet, and zebrafish embryonic stages and adult tissues used in the study.

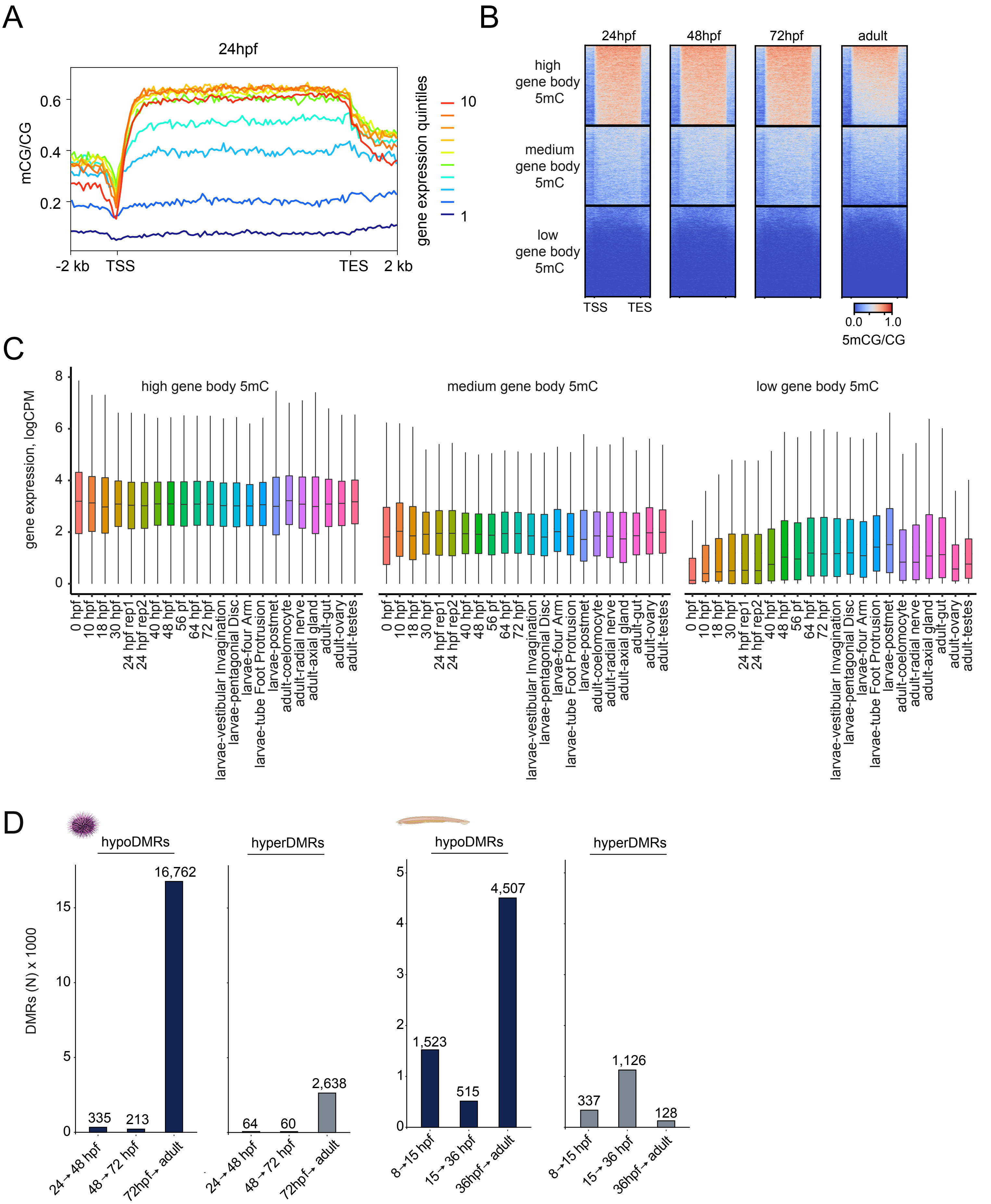


**Supplementary Figure 2. Relationships between gene body DNA methylation and gene expression in the sea urchin and the identification of differentially methylated regions. (A)** Average gene body DNA methylation of genes ranked by expression levels (10 quantiles) in 24hpf sea urchin embryos. **(B)** Heatmaps showing distinct patterns of gene body DNA methylation in sea urchin embryos and adult tissues. DNA methylation levels of gene bodies were binned into high, medium and low using k-means clustering (k=3) of 24hpf, 48hpf, 72hpf, and adult tube feet MethylC-seq samples. **(C)** Developmental gene expression dynamics of genes with high, medium and low gene body DNA methylation levels. **(D)** Number of identified differentially methylated regions (DMRs) in sea urchin and lancelet embryos and adult tissues.

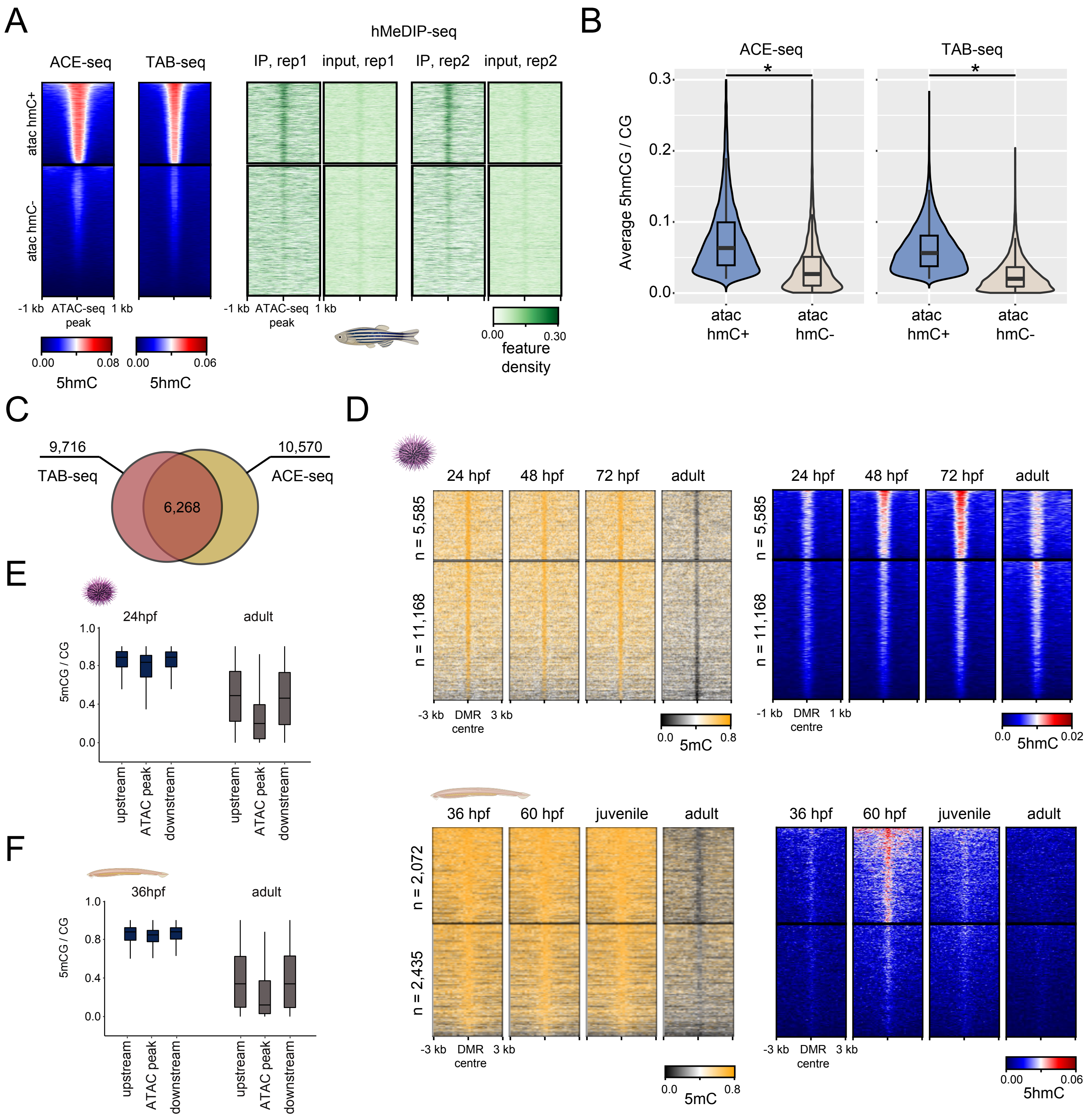


**Supplementary Figure 3. Comparison of 5hmC profiling approaches and 5mC/5hmC dynamics at DMRs**. **(A)** 5hmC enrichment as identified by ACE-seq, TAB-seq and hMeDIP-seq plotted over zebrafish ATAC-seq regions identified at 24hpf (data from Bogdanovic et al, 2016; *Nature Genetics*). K-means clustering (k=2) was performed to identify ATAC-seq regions with and without 5hmC (ATAC hmC+ and ATAC hmC-, respectively). **(B)** Average 5hmC levels within ATAC hmC+ and ATAC hmC- regions (ATAC-seq peak summit +/-250 bp). Wilcoxon rank-sum test was performed to compare average 5hmC at ATAC hmC+ and ATAC hmC- regions. **(C)** Overlap between ATAC hmC+ regions identified by ACE-seq (av. 5hmC > 5%) and ATAC hmC+ regions identified by TAB-seq (av. 5hmC > 5%). **(D)** Heatmaps showing DNA methylation (MethylC-seq) and hydroxymethylation dynamics (ACE-seq) at embryo-adult DMRs in sea urchin and lancelet. K-means clustering of ACE-data. **(E-F)** Quantification of developmental DNA methylation loss at ATAC-seq peaks corresponding to cluster 1 (Fig. 3E, F), and surrounding regions (500 bp upstream and downstream) in sea urchin **(E)** and lancelet **(F)** embryos.

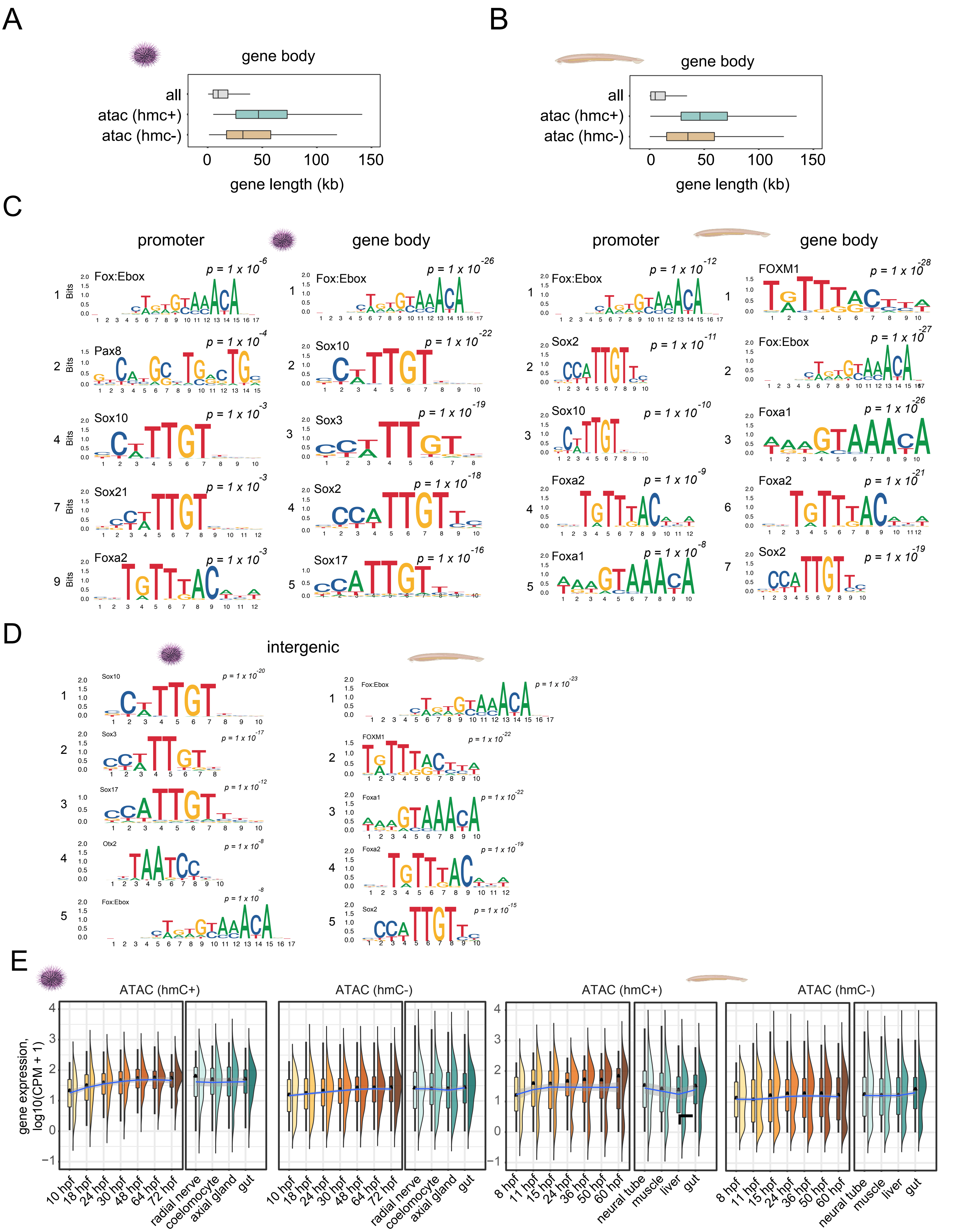


**Supplementary Figure 4. Regulatory features and gene expression profiles of 5hmC-linked genes. (A-B)** Length distributions of sea urchin **(A)** and lancelet **(B)** genes harbouring 5hmC-marked ATAC-seq peaks (gene body ATAC hmC+) and non-5hmC ATAC-seq peaks (gene body ATAC hmC-) within their gene bodies. **(C)** HOMER motif enrichment analysis of ATAC hmC+ regions associated with either gene promoters or gene bodies. **(D)** HOMER motif enrichment analysis of ATAC hmC+ regions associated with intergenic regions. **(E)** Expression profiles of gene body ATAC hmC+ genes as compared to gene body ATAC hmC- genes in sea urchin and lancelet embryos and adult tissues.

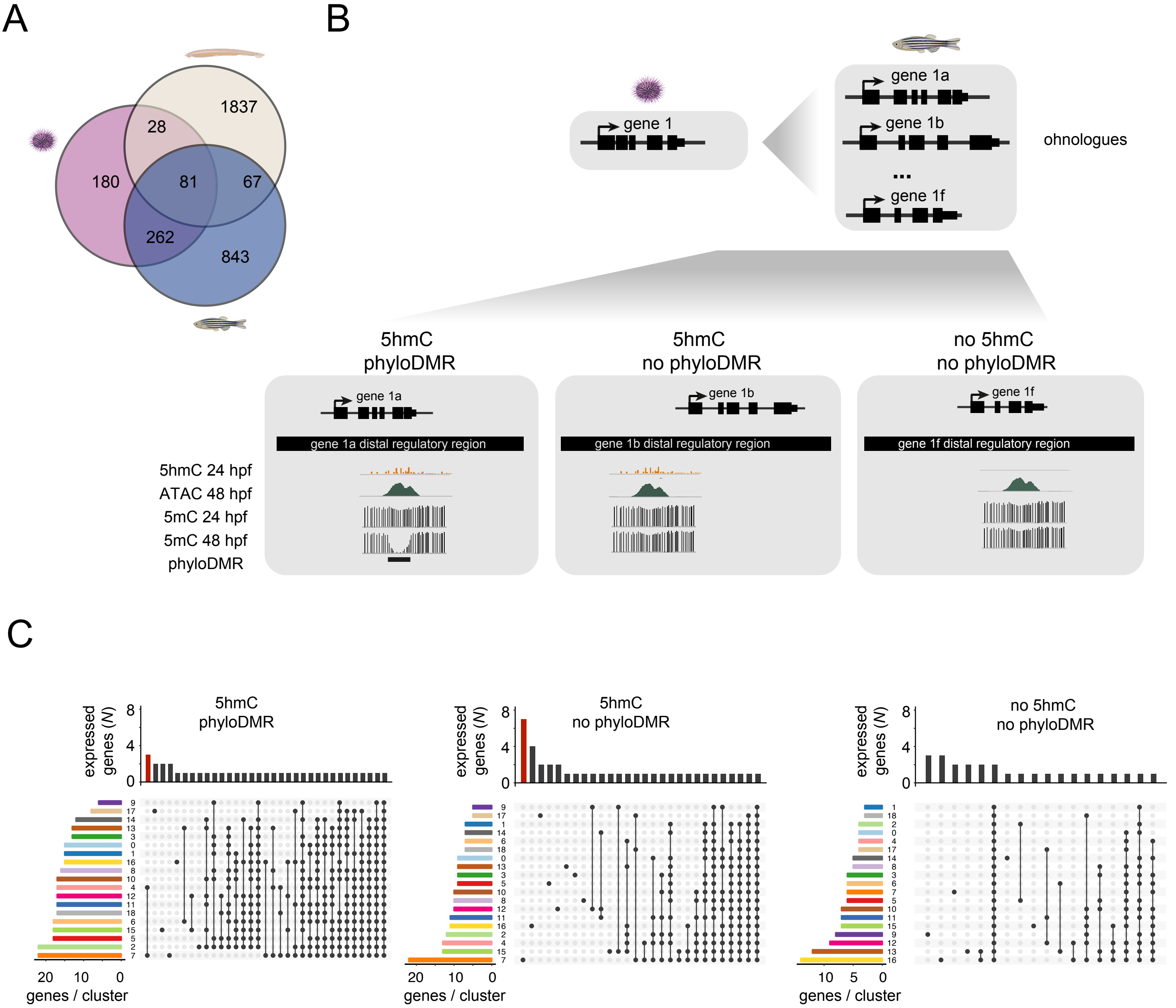


**Supplementary Figure 5. Conservation of developmental gene regulatory logic of 5hmC-marked genes in deuterostomes. (A)** Triple Venn diagram showing an overlap between ATAC hmC+ genes in sea urchin, lancelet and zebrafish. **(B)** Schematic depicting sea urchin genes associated with developmental 5hmC and their orthologous zebrafish genes, retained after three rounds of whole genome duplication (2R/3R-ohnologues). Zebrafish gene 2R/3R-ohnologues were separated into three groups: genes associated with: (i) 5hmC-marked ATAC-seq peaks overlapping phylo-DMRs (5hmC/phyloDMR), (ii) 5hmC-marked ATAC-seq peaks not overlapping phylo-DMRs (5hmC/no phyloDMR) and (iii) non-5hmC ATAC-seq peaks not overlapping phylo-DMRs (no 5hmC/no phyloDMR) in their distal regulatory domains. Distal regulatory domains were defined using GREAT gene regulatory domain definition (McLean et al, 2010). **(B)** Upset plots showing the number of genes expressed exclusively in a particular cell cluster or an intersection of clusters. A gene was considered to be expressed in a given cell cluster if it was expressed in minimum 25% of cells in the cluster. Horizontal bars depict the number of genes expressed in each cluster. Neuronal genes are marked in red.
